# Repeated signatures of balancing selection in small and large populations of guppies (*Poecilia reticulata*)

**DOI:** 10.1101/2025.06.12.659363

**Authors:** Josephine R Paris, James R Whiting, Joan Ferrer Obiol, Kimberly A Hughes, Bonnie A Fraser

## Abstract

Balancing selection is a powerful evolutionary force that maintains adaptive genetic and phenotypic diversity. Although methods to detect the footprints of balancing selection in genomic data have advanced, we still lack a clear understanding of how repeatable these signatures appear in wild populations, and how this repeatability is shaped by demographic history and existing genetic variation. The Trinidadian guppy (*Poecilia reticulata*) provides an ideal model to test the repeatability of balancing selection in the wild as there is strong evidence that negative frequency-dependent selection (NFDS) maintains colour polymorphism. Analysing whole-genome sequencing data from 11 guppy populations (*n* = 195) with contrasting demographic contexts, we apply scans of balancing selection to explore which genomic regions show evidence of repeatability. We find that populations with small *N_e_* show less genetic repeatability but still exhibit population-specific regions of elevated diversity, implicating independent balancing selection or other evolutionary mechanisms. Despite this, we identify 23 regions with repeated signatures of balancing selection, including a region on LG22 containing genes involved in colour, vision, mate choice, and social behaviour. Investigating the repeatability of balancing selection in small and large populations improves our knowledge of how demographic factors interact with selective processes to shape natural variation.

## 1. Introduction

The persistence of high polymorphism in ecologically important traits poses compelling questions in evolutionary biology. Among the mechanisms capable of maintaining balanced polymorphisms, negative frequency-dependent selection (NFDS) - whereby the fitness of a given allele increases as it becomes rarer in the population - is theoretically well-supported (1–4) and empirical examples of its prevalence continue to grow (see this issue). Recent advances in statistical approaches for identifying intermediate-frequency variants have facilitated the development of genome-wide scans for balancing selection (5,6). However, little is known about our ability to detect the signatures of balancing selection in natural populations that differ in demographic history and genetic variation. Such investigations would enhance our understanding of both the strength and constraints of NFDS, as well as its interplay with demography in shaping natural variation.

The effectiveness of balancing selection in maintaining polymorphism is dependent on population size (*N_e_*) (7,8). When *N_e_* is high, even weak balancing selection can maintain allelic diversity over extended periods of time (*N_e_*s≫1). In contrast, in small or recently bottlenecked populations, drift is expected to dominate (*N_e_*s≪1), making it difficult for selection to preserve intermediate-frequency alleles. Despite these expectations, a number of empirical studies have identified balanced polymorphisms when *N_e_* is severely reduced, particularly at immunity-related loci (e.g. (9–13), suggesting that strong balancing selection can, in some cases, overcome the effects of drift (14).

However, intermediate allele frequencies can also be produced by incomplete selective sweeps and recent population bottlenecks (6,15–17), making it difficult to distinguish true cases of balancing selection at individual loci. Population structure and admixture have also been shown to mimic the signatures of balancing selection, leading to false positives (18,19). One promising way to overcome these limitations is to search for repeated signatures of balancing selection across independent populations. The repeated maintenance of polymorphisms provides stronger evidence for selection than patterns observed within any single population, as drift and neutral mutations are unlikely to affect the same regions of the genome (20). This is especially true in populations with small *N_e_*, where the influence of drift is expected to be strong; in such cases, examining multiple populations can help distinguish true signals of selection from the background noise of neutral demographic processes. Yet despite its potential, often only single populations or species are examined for balancing selection (but see (21,22). Identifying genomic signatures of balancing selection across both small and large populations is therefore key to understanding how demographic factors interact with selective processes to shape natural variation.

Natural populations of the Trinidadian guppy (*Poecilia reticulata*) provide an ideal system to investigate the repeatability of genomic signatures of balancing selection. At the phenotypic level there is strong, repeatable evidence for NFDS. Male guppies have highly variable and heritable colour patterns, with levels of genetic variance that exceed expectations under recurrent mutation and purifying selection alone, implicating balancing selection (23). NFDS, specifically, has been demonstrated for colour pattern frequencies in natural populations. In a series of independent experiments, males with colour patterns that were manipulated to be locally rare had a significant reproductive advantage (24–26) and significant survival advantage compared to common patterns (27,28). It is likely that female mate choice is the primary driver of maintaining colour polymorphism; indeed, female preference for rare colour patterns has been shown to be widespread across multiple populations (29,30). In addition to strong evidence for NFDS, population structure and demography is well documented in the Northern Trinidad guppy system (31–34).

Here, the northern mountain range isolates populations among rivers and waterfalls, resulting in upstream populations founded by small numbers, likely from their downstream counterparts (35,36). Finally, experimentally introduced guppy populations, where downstream ‘high-predation’ guppies are transplanted to upstream guppy, and predator-free, locations (37–40), gives a unique opportunity to examine populations with known ages and sizes of founding bottleneck events (41–43). Taken together, the guppy system offers a powerful comparative framework, integrating natural populations and experimental introductions, with contrasting demographic histories and well-characterised ecological drivers and phenotypic responses to NFDS. Here, we perform genome-wide scans across 11 guppy populations to test for the repeatability of balancing selection. We selected populations with empirically documented ecological drivers of NFDS - specifically, female preference for rare male phenotypes (29), which also span a range of contrasting demographic histories (32,34). First, we quantify population structure, demography, divergence, and genetic diversity to categorise populations into small and large demographic groups. We then examine regions of elevated genetic diversity found exclusively in small populations and assess their repeatability using change-point detection methods. Next, we apply newly developed model-based approaches (44–47) to scan for balancing selection, to identify whether, and where, we find repeatable signatures across populations. Finally, we examine candidate loci that are consistently found to be under balancing selection to assess their potential links to phenotypic traits targeted by NFDS.

## 2. Methods

### (a) Study sites and sample collection

Guppy samples (n = 195) were collected from 11 sites (across three drainages) in the Northern mountain ranges of Trinidad (figure 1a), ranging from 10 to 20 samples per site (table S1). For the Guanapo (Caroni drainage), Aripo (Caroni drainage) and Marianne (Northern drainage) rivers, guppies were sampled both downstream and upstream of waterfalls. Downstream (D) populations are typified by high-predation pressure (48) and are considered to be the source of naturally-colonised upstream populations. Naturally-colonised upstream populations (U) are hypothesised to have been founded by a small number of individuals, and are generally typified by lower genetic diversity and low predation pressure (*i.e*., “low-predation” populations) (32,33). We also sampled experimentally-introduced populations from the El Cedro (Caroni drainage) and Turure (Oropouche drainage) rivers and their downstream counterparts. In 1981, ~100 guppies were introduced from a downstream El Cedro site into a guppy-predator-free upstream site in the same tributary (37,38). In 1956, ~200 guppies were introduced from a downstream Guanapo site into an upstream site in the Turure river (40), where they subsequently naturally colonised a previously occupied downstream Turure site (49,50). All samples were stored in either 95% EtOH at −20° C or RNeasy at 4° C prior to DNA extraction.

**Figure 1.**
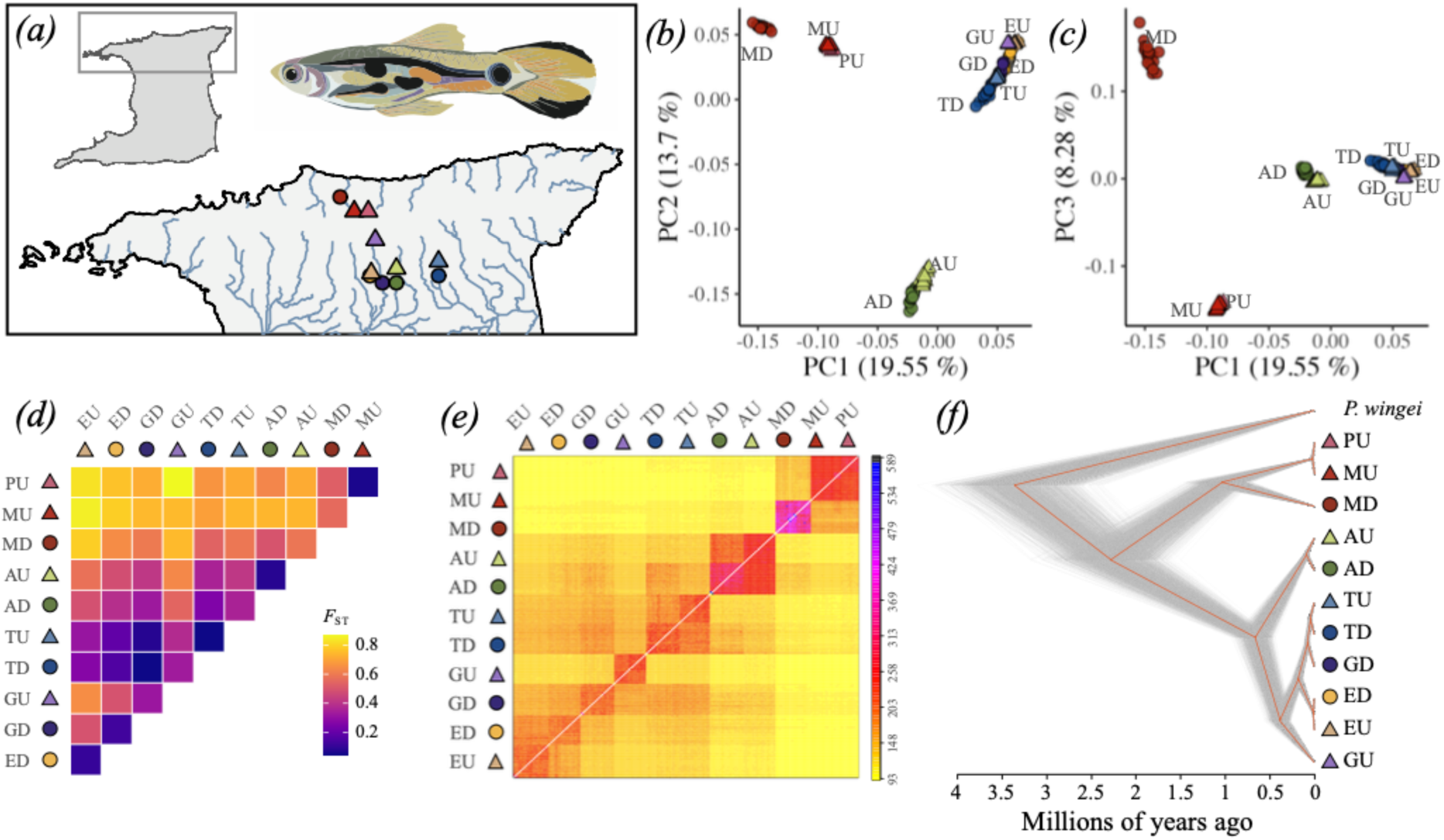
Overview of the guppy study system and analysis of population structure and divergence. (*a*) Location of the 11 study sites in the Northern Mountain ranges of Trinidad. Inset shows the location (black box) of the main map of the island of Trinidad. Populations are colour-coded by river: Guanapo (purple); El Cedro (orange); Aripo (green); Turure (blue); Marianne (red); Paria (pink). Downstream (D) populations are denoted with a circle. Upstream (U) populations are denoted by a triangle. (*b*) Principal Components Analysis (PCA) showing PC1 (~20%) and PC2 (~14%). (*c*) PCA showing PC1 (~20%) and PC3 (~8%). (*d*) Heatmap of pairwise *F*ST calculated between all 11 populations, where purple represents lower *F*ST and yellow represents higher *F*ST. (*e*) Pairwise co-ancestry matrix between individuals, with light yellow indicating low co-ancestry, and darker yellows, oranges and reds to pink indicating progressively higher co-ancestry. (*f*) Dated divergence tree highlighting the relationships and divergence between populations. Guppy picture © by Erica Robertson.

### (b) DNA extraction, whole-genome sequencing, and variant calling

Total genomic DNA was extracted using the Qiagen DNeasy Blood and Tissue kit (QIAGEN, Germany), including an on-column RNAse step. Low Input Transposase Enabled (LITE) DNA libraries were prepared at The Earlham Institute, UK. Libraries were sequenced on an Illumina HiSeq 4000 (150bp paired-end) with a target sequencing coverage of 10x per sample. Sequencing statistics can be found in table S2.

Variant calling followed previous approaches (33,41). Briefly, reads were cleaned using cutadapt (51), aligned to the guppy genome (GCA_904066995.1; (52) using bwa mem v0.7.17 (53), and sorted and indexed using samtools v1.17 (54). Quality control and coverage of samples were checked using samtools *stat, flagstat* and *idxstats*. Variant calling was performed using GATK4 v4.1.8.1 (55). Variants were recalibrated using a truth-set generated from high-coverage, PCR-free sequencing data (52). Variants were filtered using quality control filters (QD < 2.0 | FS > 60.0 | MQ < 40.0 | HaplotypeScore > 13.0), retaining only biallelic SNPs, at a minimum depth of 4 and a maximum depth of 200. Population-specific VCF files were created and filtered at either a maximum missing value of 50% or 80% using vcftools (56) and then re-combined. For analyses requiring phased variants, phasing was performed using Beagle v2 (57), followed by shapeit v2 (58). To incorporate recently documented genome rearrangements (on LG12 and LG20) (33,59,60), output file coordinates were lifted over using custom scripts. Further details on the specific variant filtering performed for each analysis can be found in table S3.

For divergence time estimates and for polarising ancestral alleles, we also made use of data for two other species: *Poecilia wingei* (sister) and *Poecilia picta* (outgroup). *P. wingei* WGS data (n=6) were accessed via the European Nucleotide Archive (PRJEB26489) (61). For *P. picta*, we performed WGS on ten laboratory-reared individuals (5 males and 5 females). Data were sequenced on an Illumina HiSeq 2500 (150bp paired-end; table S2).

### (c) Population structure, genetic diversity, and demographic history

To assess population structure and shared ancestry, we performed a Principal Components Analysis (PCA) in Plink v.19 (62), estimated mean pairwise *F_ST_* in PopGenome v2.75 (63), and used fineStructure v4.1.1 (64) to build a co-ancestry matrix based on haplotype relationships.

For divergence dating, we used a multispecies coalescent (MSC) approach implemented in SNAPPER (65). We followed the recommendations of Stange et al. (66) by constraining the root of the tree (divergence between *P. reticulata* and *P. wingei*) to follow a normal distribution with a mean of 3.41 mya and a standard deviation of 0.329 (33) (supplementary methods S1). Relative cross-coalescence rates between upstream-downstream pairs and contemporary *N_e_* for each population were estimated using ancestral recombination graphs (ARGs) in Relate v1.1.6 (67,68). Analysis was performed using a starting effective population size of 1×10^4^, a mutation rate of 2.90 × 10^−9^ per site per generation (69), and smoothed recombination maps for each chromosome (70). The ancestral allele was assigned as the one more common in the sister (*P. wingei*) and outgroup (*P. picta*). To convert the ARGs into estimates of population size, we used the script EstimatePopulationSize.sh using a generation time of 7.9 months (0.66 years) (71).

Nucleotide diversity (π) and Tajima’s *D* were calculated in 50 Kb non-overlapping windows in PopGenome v2.75 (63). Observed heterozygosity (*H*_o_) was estimated per site per population in vcftools v0.1.16 (56) using the --hardy parameter and was summarised as a mean in 50 Kb windows. Change point detection (CPD) was performed on the Tajima’s *D* estimates per population using the changepoint package (72) in R, using the cpt.mean function with the BinSeg method, an SIC penalty and a Q of 100. Repeated CPD regions with elevated Tajima’s *D* were investigated further using lostruct (73) by performing local PCA in windows of 100 SNPs, and by calculating LD using the –r2 square option in Plink, plotting the results with LDheatmap (74).

### (d) Scanning the genome for balancing selection

Scans for balancing selection were conducted using BalLeRMix+ (44,45), which implements a series of composite-likelihood ratio (CLR) statistics (known as “*B* statistics”) by modelling local deviations in the allele frequency spectrum as it is locally perturbed by selection. The CLR test includes the flanking region surrounding the site of interest, avoiding issues of window size selection. We chose BalLeRMix+ over other scans of balancing selection because it has been shown to be robust under different demographic scenarios (population expansions and bottlenecks), different strengths and ages of selection, and window-size selection and recombination rate biases (44). Amongst the five *B* statistics, the *B_2_* statistics, which model the allele frequency spectrum whilst incorporating substitution information, have been shown to be the most powerful in detecting balancing selection (44).

We used BalLeRMix+ to calculate *B_2_* statistics using both the derived allele frequencies (DAF; supplementary methods S2) and the minor allele frequencies (MAF), inferred for each population. Prior to calculating allele frequencies, the VCF file was filtered to account for potential artefacts which might lead to false positives, including mappability, repeat-rich regions, and high/low coverage regions (supplementary methods S3). We compared the *B_2_* DAF and *B_2_* MAF statistics due to some uncertainty in our identification of the derived allele in some of the populations (figure S1 & figure S2).

BalLeRMix+ was run using a constant recombination rate of 2.12e-06, estimated from smoothed linkage maps (70). While this constant recombination rate is likely not reflective of the entire genome, BalLeRMix+ is robust to errors in recombination rate (44). To be able to compare the BalLeRMix+ estimates with our other estimates of population genetic statistics, we summarised the *B_2_* statistic in 50 Kb windows, taking the largest CLR (which is appropriate given that the test accounts for flanking regions) and corresponding log_10_ŝ score. We excluded windows that had fewer than five variable sites. To ensure robustness of our inferences from BalLeRMix+, we also independently calculated the *β* statistic using BetaScan2 (46,47). Concordant results between the two approaches were used to increase confidence in candidate loci.

### (e) Identifying repeatable signatures of balancing selection

To identify genomic regions with repeatable signals of balancing selection, we first examined the overlap of windows in the top 1% of CLR scores across populations, but found limited consistency (see Results). While informative, such an approach is conservative, as it only considers the most extreme outliers and moreover, depends on choosing an arbitrary cutoff for significance. To overcome these limitations, we used PicMin (75), a method that evaluates whether multiple populations show a coordinated shift toward extreme values. By leveraging order statistics, PicMin offers a robust and sensitive framework for detecting repeatable signals of selection without relying on predefined significance thresholds. We used 10 million replicates for the null distribution. Significant repeatable windows were examined if they occurred in ≥ 9 populations (config_est parameter) at a −log_10_(FDR) > 4. Candidate windows were then examined further by quantifying the CLR statistic and log_10_ŝ estimates (where ŝ > 0 indicates balancing selection and ŝ < 0 implies positive selection). To explore the genes in candidate regions, we performed a new annotation of the guppy genome using BRAKER3 (76) (supplementary methods S4).

## 3. Results

### (a) Genetic structure and divergence reveals strong drainage effects and traces of introduction history

Assessment of population structure, co-ancestry, and divergence among the 195 guppies sampled from 11 sites across Northern Trinidad revealed strong genetic clustering that reflects the river drainages and the known introduction history of the experimental populations (figure 1; figure S3). PC1 (19.6%) separated populations from the Northern drainage populations (MD, MU, PU) from those in the Caroni drainage, while PC2 (13.7%) distinguished the Aripo populations (AD, AU), and PC3 (8.3%) further separated MD from its upstream counterparts (MU, PU), which clustered more closely together (figure 1b, 1c).

Pairwise F*_ST_* estimates among natural downstream-upstream pairs ranged from low (0.08 between AD and AU), modest (0.32 between GD and GU) to high (0.56 between MD and MU), indicating varying degrees of differentiation due to the time since upstream colonisation and subsequent gene flow (figure 1d). Experimentally-introduced populations reflected their sources to varying extents: EU was moderately diverged from ED (F*_ST_* = 0.10), while TU was genetically proximate to both its source (GD; F*_ST_* = 0.07) and its downstream counterpart (TD; F*_ST_* = 0.04), indicative of introduction history and post-introduction gene flow, respectively.

Coancestry analyses showed a clear clustering by drainage, strong similarity between MU and PU, and a very low shared co-ancestry of GU with all other populations, even with its putative source (GD) (figure 1e).

Divergence time estimates inferred deep splits between the Northern and Caroni drainages (~2.2 mya), and more recent divergences among introduced and source populations. However, the latter were older than known introduction times, most likely due to biases from the MSC model assumption of no post-divergence gene flow (figure 1f). However, relative cross-coalescence rates (RCCR) (which unlike divergence dating, incorporate gene flow) showed expected shallow splits between introduction pairs (e.g. ED–EU, TD–TU), more recent divergence between GD–GU, and deeper divergence for Northern drainage pairs (MD–MU, MD–PU) (figure S4). Together, these results point to strong drainage-driven structure and a mosaic of both shared and independent demographic histories across the 11 guppy populations.

### (b) Variation in *Ne* and genetic diversity across upstream and downstream guppy populations

Genome-wide estimates of diversity demonstrated that guppies occupying downstream (D) sites, generally harbour more genetic variation compared to upstream (U) sites (table 1). Four of the six upstream sites showed much lower genetic diversity (GU, MU, PU, EU), with two showing extremely reduced diversity (GU and MU; figure 2). The upstream populations, AU and TU, did not show as much reduced diversity, likely due to higher amounts of connectivity with their downstream counterparts (figure S4). Estimates of *N_e_* further highlighted the low diversity of these four upstream populations, in particular GU and MU (table 1; figure S5). Of the experimentally-introduced sites, TU showed similar amounts of diversity and a similar *N_e_* relative to its source population (GD), but EU had a lower diversity and *N_e_* compared to its source (ED), and ED had the lowest diversity and *N_e_* compared to other downstream populations. Together, these analyses allowed us to classify the populations into small populations: GU, MU, PU and EU; and large populations: GD, AD, AU, TD, TU, MD, ED.

**Figure 2.**
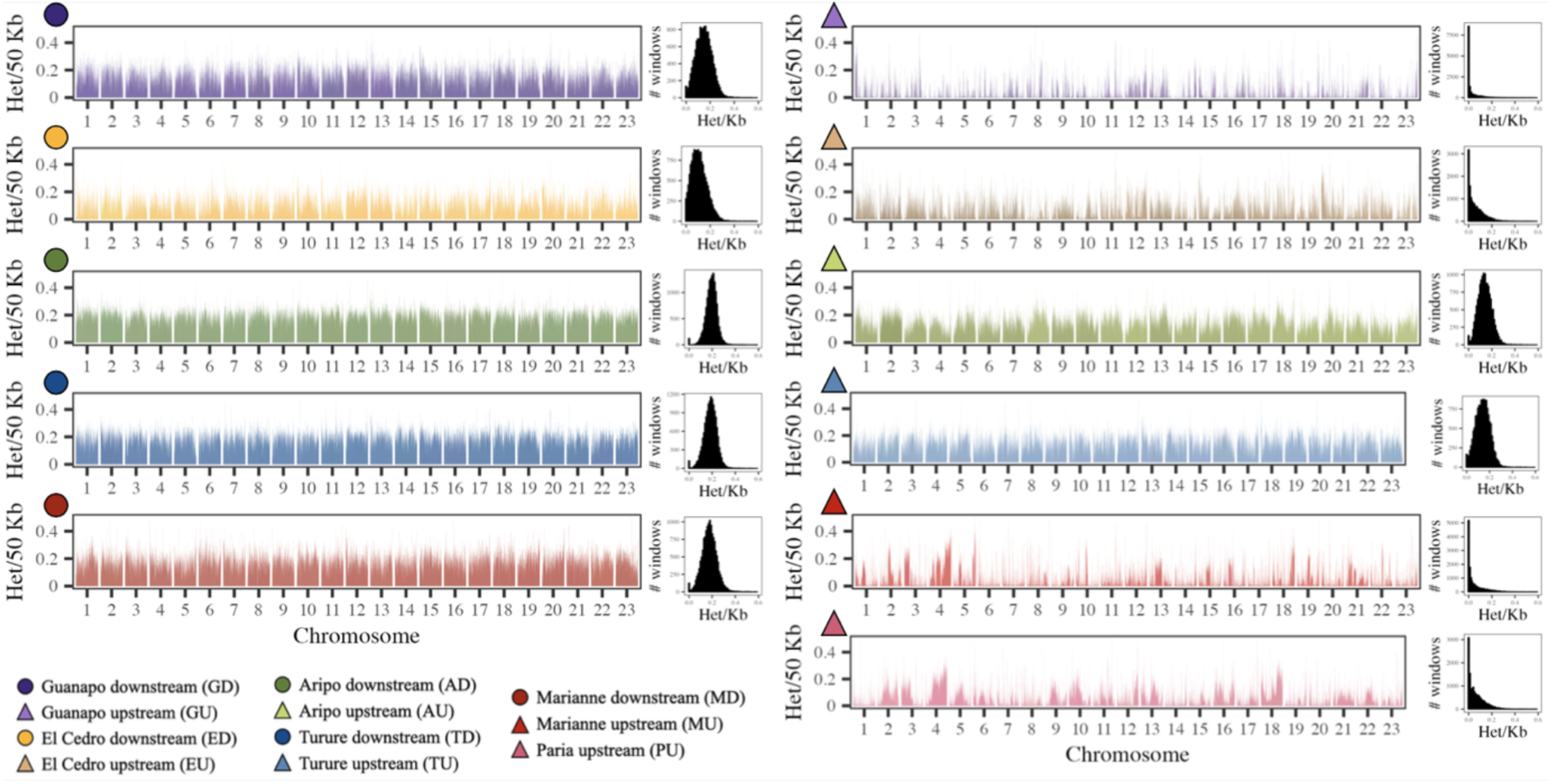
Heterozygosity estimated across the 23 chromosomes of the guppy genome for the 11 studied populations. Plots represent *H*o estimated in 50 Kb windows across the 23 chromosomes of the guppy genome. Plots on the right summarise the density distributions. Downstream populations (denoted by circles) generally show higher levels of genetic diversity than upstream populations (denoted with triangles). Rivers are colour-coded: Guanapo (purple); El Cedro (orange); Aripo (green); Turure (blue); Marianne (red); Paria (pink).

**Table 1.** Summary of effective population size (*N_e_*) and metrics of genetic diversity calculated for each of the 11 guppy populations.

| Population | Code | Ne | Heterozygosity | Nucleotide diversity ($\pi$) |
| --- | --- | --- | --- | --- |
| Guanapo Downstream | GD | 748 | 0.1467 | 0.0007 |
| Guanpao Upstream | GU | 52 | 0.0017 | 0.0000 |
| El Cedro Downstream | ED | 357 | 0.1033 | 0.0005 |
| El Cedro Upstream | EU | 121 | 0.0419 | 0.0002 |
| Aripo Downstream | AD | 840 | 0.1961 | 0.0010 |
| Aripo Upstream | AU | 271 | 0.1374 | 0.0007 |
| Turure Downstream | TD | 1422 | 0.1866 | 0.0009 |
| Turure Upstream | TU | 724 | 0.1403 | 0.0007 |
| Marianne Downstream | MD | 399 | 0.1752 | 0.0008 |
| Marianne Upstream | MU | 120 | 0.0176 | 0.0001 |
| Paria Upstream | PU | 208 | 0.0445 | 0.0002 |

### (c) Distinctive diversity patterns and shared Tajima’s *D* outliers in small populations

We found that several small populations exhibited a characteristic “sawtooth” pattern of diversity, marked by distinct islands of elevated diversity (figure 2). Furthermore, the distribution of Tajima’s *D* values revealed greater variance in small populations compared to large ones, characterised by a distinctly bimodal pattern. Distributions in small populations were skewed toward both highly negative values (*i.e.*, an excess of rare alleles), and highly positive values (*i.e.*, an excess of intermediate-frequency alleles; figure S6). To explore this further, we performed change point detection (CPD) on Tajima’s *D* (table S4), finding that the small populations GU, EU, and to a lesser extent, MU, have more and longer segments of consistently elevated Tajima’s *D*. Such patterns were not identified in any other populations. To investigate shared regions of elevated diversity in these three small populations, we examined potential overlaps in CPD regions. While numerous pairwise overlaps were detected when Tajima’s *D* = 2-3, only one region was shared across all three populations (LG18: 25.05 - 25.65 Mb). In the more extreme Tajima’s *D* CPD regions (>3), we identified a shared region between MU and GU on LG3 (7.65 - 7.80 Mb), and another between EU and MU on LG22 (12.25 - 13.35 Mb). Further investigation of these three regions showed that they did not share a common recombination landscape (inferred using local PCA; figure S7), but showed elevated LD in nearly all populations (figure S8).

### (d) Balancing selection signatures show limited repeatability across populations

Using the *B_2_* statistic, we found signatures of selection across the 11 populations using both the derived allele frequency (DAF) spectra (table S5) and the minor allele frequency (MAF) spectra (table S6). Outlier windows (defined as ≥ 99% of the *B_2_* CLR distribution) differed among populations, with no obvious regions showing high repeatability (figure S9 & S10). The large majority of outlier windows identified as being under strong balancing selection using the MAF spectra were also identified as outliers using the DAF spectra (table S7). We therefore used the results obtained via modelling of the DAF spectra for the remainder of our analyses.

The majority of outlier windows in each population showed evidence of balancing selection (ŝ > 0). Only the introduced large TU population showed a large fraction of windows showing positive selection (where ŝ is −3), and noticeably, none of the small populations (GU, EU, MU, PU) had any windows with a signal of positive selection (table S5). This is expected as *B_2_* models the global SFS to the local SFS and many of the small populations had a large amount of fixed variation (figure 2). *B_2_* windows showing the strongest signals of balancing selection (≥ 99% and ŝ = 9) exhibited limited, but some overlap with candidate windows (≥ 99%) identified by BetaScan (table S8). These *B_2_* windows were mostly located in the right-hand tails of the Tajima’s *D* and *π* distributions (figure S11). However, repeatability in these windows across populations was largely inconsistent, with no outlier windows shared in eight or more populations, only one window found in seven populations, and just seven windows found in five populations (table S9).

To further explore potential repeatability in signatures of balancing selection among our populations, we used PicMin, hypothesising that this would increase our power to detect candidate windows under balancing selection. Using PicMin, we identified 43 windows with a −log_10_(FDR) > 4 supported by ≥ 9 populations (N=16 windows in 9 populations; N=18 windows in 10 populations; N=9 windows in 11 populations; table S10). Of these, 23 windows (found across 13 chromosomes) had an ŝ>0 in all 11 populations and therefore show repeatability only for balancing selection (figure 3).

**Figure 3.**
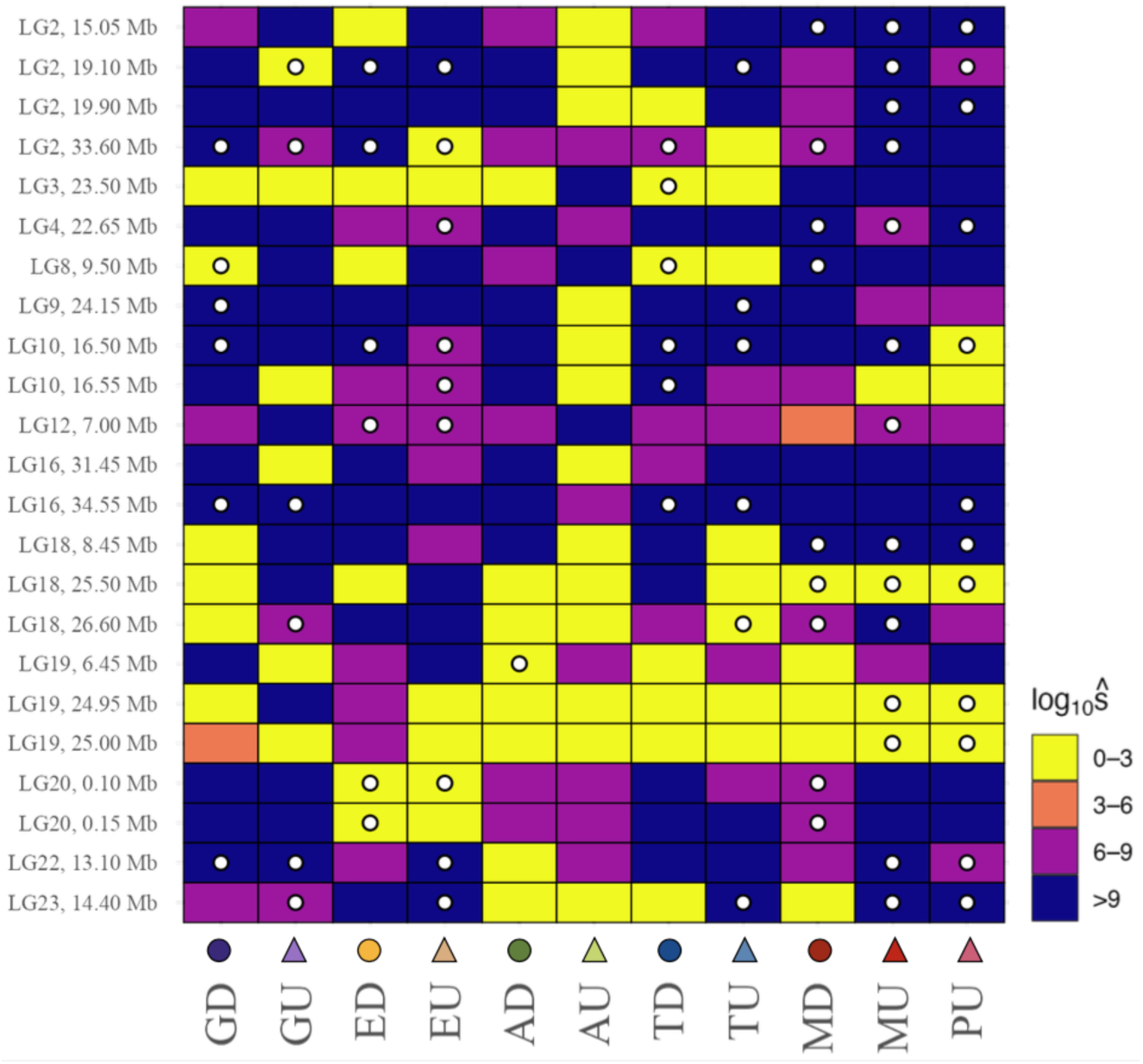
Heatmap showing the 23 candidate windows and their patterns of repeatability across the 11 guppy populations. Windows are numerically ordered by chromosome and window on the y-axis and the populations are ordered along the x-axis. Squares are coloured by their signature of balancing selection (log_10_ŝ) as indicated in the legend. White circles within each square denote significant windows (*i.e.*, top 1% of *B_2_* scores) for any given population.

### (e) Balancing selection candidates

We investigated the 23 candidate regions with the strongest evidence for repeated balancing selection as identified using PicMin, by integrating population-level *B_2_* scores (≥ 99%), *β* statistics (>90%), and functional gene annotations (table S11). Several regions emerged as particularly interesting due to their high statistical support and the presence of genes with functional annotations potentially related to NFDS in guppies (see Discussion). For example, a region on LG2 (~33.6 Mb) was among the top 1% of *β* scores in all 11 populations and among the top 1% of *B_2_* scores in seven populations. On LG10 (~16.5 Mb), two adjacent windows showed strong evidence of balancing selection, with the first window in the top 1% of both *B_2_* and *β* scores across seven populations. A region on LG16 (~34.6 Mb) was in the top 1% of *B_2_* scores in five populations and in the top 5% of *β* scores in six populations. On LG18 (~26.6 Mb), we identified a region containing seven genes which scored in the top 1% of *B_2_* scores in four populations and in the top 1% of *β* scores in seven populations. Finally, a window on LG22 (~13.10 Mb) containing five genes (figure 4), was in the top 1% of *B_2_* scores in five populations and the top 5% of *β* scores in six populations. Notably, two of these regions, on LG18 and LG22, also overlapped with CPD segments showing elevated Tajima’s *D* in the small populations, further supporting their role as repeated targets of balancing selection.

**Figure 4.**
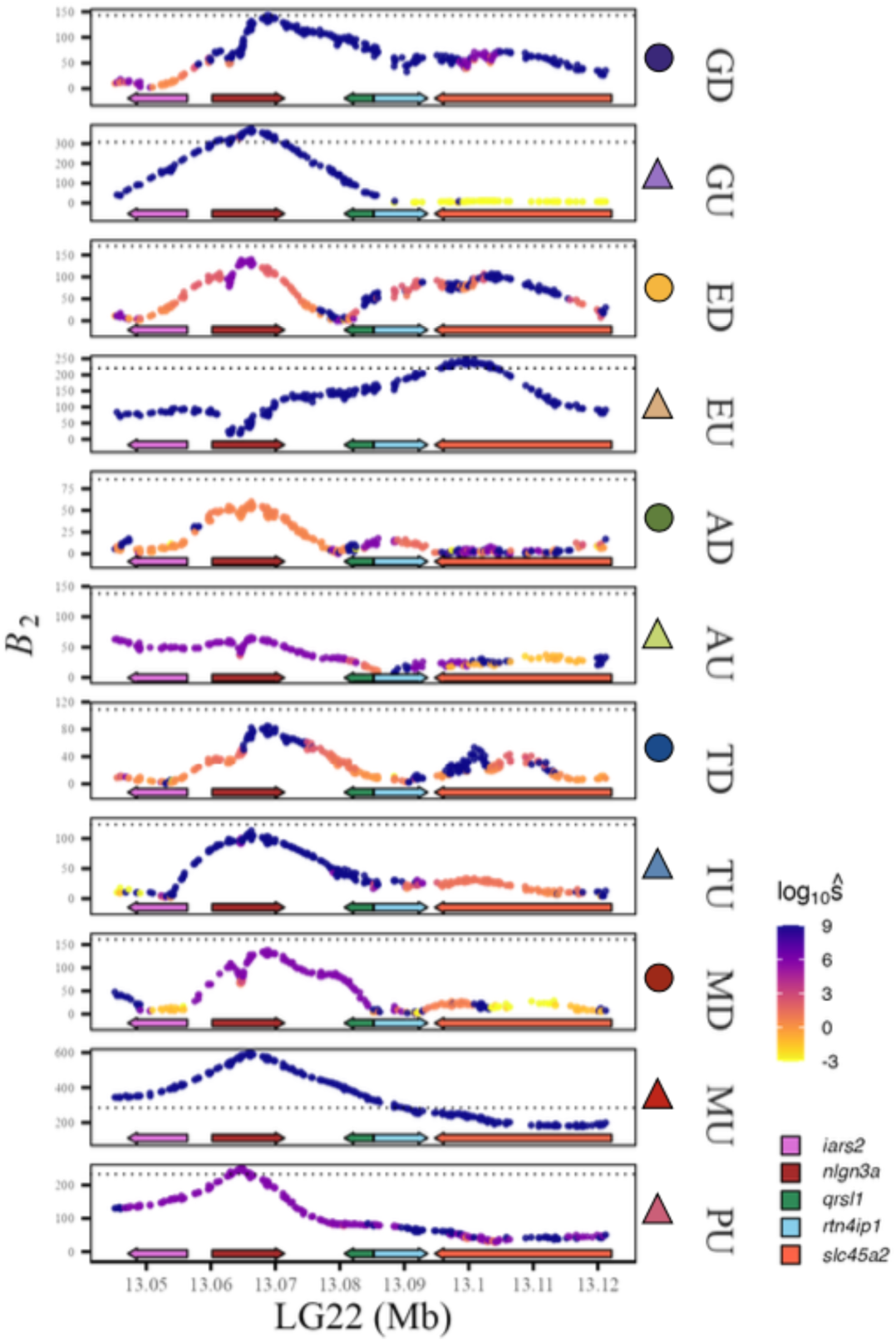
Evidence for repeated balancing selection across 11 guppy populations on LG22 (13.03 - 13.1 Mb). *B*_2_ scores across the genomic region on chromosome 22 surrounding four genes, from left to right: *iars2* (*isoleucyl-tRNA synthetase 2, mitochondrial*; in pink); *nlgn3a* (*neuroligin 3a*, in brown); *qrsl1* (*glutaminyl-tRNA amidotransferase subunit*; in green); *rtn4ip1* (*reticulon 4 interacting protein 1*; in blue); *slc45a2* (*solute carrier family 45 member 2*; in orange). Population *B*_2_ scores are coloured by their signature of selection (log_10_ŝ) as indicated in the legend.

We explored the candidate region on LG22 in more detail as it was among the top 1% of *B_2_* scores for five populations, including all four of the small populations, where log_10_ŝ was also high: GU ŝ = 9, EU ŝ = 9, MU ŝ = 9, PU ŝ = 6. The LG22 window also overlapped with the region identified in the Tajima’s *D* CPD analysis (LG22: 12.25 - 13.35 Mb) and contained genes relevant to the guppy NFDS phenotype (see Discussion; figure 4). With the exception of the introduced and small EU population, all populations showed elevated *B₂* over the *nlgn3a* gene. Signals across other genes in the window varied by population: GD, ED, AD and TU maintained high *B₂* scores across both *qrsl1* and *rtn4ip1*, while EU exhibited even higher B₂ over these genes compared to *nlgn3a* and other populations. Increased *B₂* scores over *slc45a2* were also observed in GD, ED, EU, AU, TD, TU and MD. Examination of per-SNP *β* scores within the region revealed a pronounced peak in all populations (in the top 1% in eight of them), again with the exception of EU (figure S12). LD patterns in the candidate region revealed strong and extensive LD in the small populations (GU, EU, MU, PU) consistent with reduced recombination or long-range haplotype maintenance (figure S13). In contrast, larger populations exhibited more fragmented LD, but a region of elevated LD was apparent surrounding the coordinates encompassing the gene *nlgn3a*.

## 4. Discussion

Despite its potential to maintain stable polymorphisms, the genomic signatures of NFDS, and balancing selection more broadly, remain understudied in natural populations, limiting our understanding of the strength and constraints of NFDS in shaping natural variation. We analysed 11 guppy populations spanning a range of demographic contexts to investigate the repeatability of balancing selection. Populations exhibiting reduced diversity and low *N_e_* were found to harbour localised regions of elevated diversity, but the repeatability of these regions across populations was low. Using new statistical methods to scan the genome for balancing selection, we initially found limited overlap in the top 1% of outlier regions across all populations. However, by taking advantage of the repeatability of NFDS in this system, we applied an order-based statistical framework, identifying 23 genomic regions (across 13 chromosomes) with consistent signatures of selection. Further investigation of these regions revealed genes with putative links to the targets of NFDS in guppies, such as those involved in colour, vision, mate choice, and social behaviour. Together, these results suggest that, despite demographic heterogeneity, certain genomic regions are recurrently targeted by balancing selection, reflecting shared ecological drivers of NFDS, conserved adaptive functions, and evolutionary constraint on where in the genome adaptive variation can persist.

Our results provide new insights into how balancing selection operates in small, bottlenecked populations, where the effects of drift are expected to overwhelm selection. Rather than relying solely on an upstream-downstream classification, we used multiple metrics of genetic diversity and effective population size (*N_e_*) to classify the 11 populations, revealing that not all upstream populations are equally small. For example, AU and TU were found to retain diversity levels more comparable to their downstream counterparts, likely due to higher gene flow (34). In contrast, the small populations (GU, MU, PU, and EU) exhibited a distinctive “sawtooth” pattern of diversity, resembling patterns observed previously in other small populations (11,77), attributed to both recent inbreeding and balancing selection. Indeed, the bimodal distribution of Tajima’s *D*, with both strongly negative and positive values, suggests that the observed diversity peaks reflect the influence of multiple, potentially interacting evolutionary processes operating in small populations. The guppy system offers a unique opportunity to compare these islands of elevated diversity with independent signals of balancing selection, and to assess their repeatability across populations, providing insights into the mechanisms that maintain genetic diversity in small populations.

Recombination, and how it interacts with selection and drift, may be playing an important role in maintaining diversity. In small populations, genetic drift can generate LD which indirectly favours modifier alleles that increase recombination, thereby enhancing the probability that beneficial combinations of alleles arise and persist (78). Empirical evidence supports the idea that recombination rate variation can modulate the efficiency of selection in small populations (79), and inbreeding alone can cause the indirect selection of higher recombination rates (80). Moreover, regions with elevated recombination may retain more diversity even under strong drift, potentially helping to preserve intermediate-frequency alleles (81). In our small populations, elevated LD in candidate regions with consistently high Tajima’s *D*, along with the absence of shared reduced recombination patterns, suggest that recombination, and its interplay with drift and selection, may be a key, and often overlooked factor influencing how diversity is maintained in small populations. Whilst we detected regions of elevated Tajima’s *D* in the small populations, they exhibited the lowest repeatability in balancing selection signals. This raises an important question: is balancing selection genuinely rarer in small populations, or simply harder to detect? While drift undoubtedly reduces the efficacy of selection and may limit the persistence of polymorphisms, it is also possible that stochastic demographic effects, such as inbreeding, or expansions after a bottleneck, could obscure otherwise detectable signals.

Despite our documented differences in demographic contexts among the 11 populations, we identified repeated signatures of selection at 23 regions, which included candidate genes related to the NFDS phenotype. These included two copies of a sacsin-like gene, best known for its role in neuronal function (82). Mutations in this gene cause retinal abnormalities in zebrafish (83), and it has also been associated with sociality in cichlids (84). We also detected signatures of balancing selection on a region containing repeated copies of protocadherin genes, which belong to a large family of neural cell-adhesion molecules implicated in synaptic connectivity and neuronal identity (85). This suggests a potential link between NFDS and variation in neural processing of colour cues, sensory perception, or behavioural responses. Notably, cadherin genes have been previously implicated in adaptation to predation regimes in guppies (33,41). Balancing selection was also evident in a region containing two *adgrb1* (BAI1) genes which in knockout mice are associated with spatial memory deficits, altered synaptic plasticity (86), and differences in social discrimination (87). One region overlapping with elevated Tajima’s *D* on LG18 included several compelling genes: *ednrba* involved in pigmentation and expressed in melanocyte, iridophore, and xanthophore cells in zebrafish (88), two copies of *nlrp12l,* a key innate immune regulator (89); and *dclk2*, involved in dendritic and retinal development (90,91).

In the LG22 candidate region, we found particularly strong evidence for balancing selection at the *nlgn3a* (*neuroligin-3*) gene, a synaptic adhesion molecule linked to social novelty responses in mice (92–94). Moreover, expression of this gene has been specifically linked to mate preference in Poeciliids (95,96), including guppies (97,98). Downstream of *nlgn3a*, we found variable signatures of balancing selection in *qrsl1*, essential to mitochondrial protein synthesis, and *rtn4ip1*, important to optic health (99), which could influence energy expenditure related to mating displays and visual processing critical to mate recognition. Finally, we observed varied signatures of balancing and positive selection on *slc45a2*, a key gene involved in melanin synthesis (100). Future studies could validate the functional relevance of these genes by integrating gene expression analyses, CRISPR-based functional knockouts, or behavioural assays linking genotype to colour variation, visual discrimination, or social responsiveness under controlled ecological and demographic contexts.

It is likely that we missed some regions under balancing selection, as the statistical approaches we used are designed to identify excess intermediate-frequency polymorphisms, which are characteristic of long-term balancing selection. Such methods are less sensitive to detecting ancient trans-species polymorphisms (101) or recent balancing selection, such as those arising during a selective sweep (5,8). Nevertheless, long-term balancing selection is the most plausible model for our hypothesised mode of NFDS, *i.e.*, that male colour polymorphisms have been maintained across guppy populations through consistent female preference for rare male phenotypes (29,30). However, the experimentally introduced populations have only recently experienced changes in predation selection pressures, and any newly emerging balanced polymorphisms may not yet have had sufficient time to exhibit strong genomic signatures (although see (17)). Such signatures would be particularly interesting to investigate further as recent empirical evidence has shown that fixation of beneficial alleles often follows an initial period of balancing selection (*i.e.*, a staggered sweep model) (102).

We employed conservative filters to avoid false positives arising from repetitive, duplicated, or misassembled regions of the genome. While this approach reduces spurious signals (103), it also likely excluded more complex loci, including those in gene families known to evolve under balancing selection, such as immunity genes. Indeed, MHC genes are known to exist as multiple copies in guppies (104–106). Similarly, we did not detect repeatable balancing selection on male-specific regions of elevated diversity on the Y chromosome (LG12) (107), nor on the more complex haplotypes previously implicated in male colour variation (108). Future work using long-read sequencing and pangenome-based approaches will improve the detection of balancing selection in complex regions (109), especially at loci with high copy number variation and/or extensive structural variation (22,110,111).

Finally, as with most genome scan approaches, our detection framework is inherently biased toward loci of large effect (112). Traits like male colouration and mate choice in guppies are likely polygenic, meaning that weaker or more diffuse selection signals could go undetected (113). Conversely, some regions identified here may represent false positives, since patterns of intermediate allele frequency can also arise from incomplete sweeps, population structure, or admixture (18). However, our divergence tree showed no evidence for recent admixture across drainages, reducing the likelihood that our most repeatable signals are the result of introgression. Overall, our strategy of comparing populations with contrasting predation-selective histories and demographic contexts strengthens the inference that our candidate regions represent robust targets of balancing selection.

We primarily relied on the *B_2_* statistic using the derived allele frequency (DAF) spectrum (44,45), alongside the rank-based PicMin approach (75) to identify candidate regions of repeatable balancing selection. To strengthen the evidence for these candidates, we also compared them with the *B_2_* statistic modelled using the minor allele frequency (MAF) spectra, *β* statistics (46,47), and standard diversity statistics. We chose to use the *B_2_* statistic modelled on the DAF spectra because it incorporates both substitutions and polymorphisms, and has been shown to outperform other methods across a wide range of demographic scenarios and recombination rate heterogeneity (44); the latter being particularly critical as we applied a uniform recombination map. One challenge of DAF-spectra-based approaches is the requirement for reliable inference of the ancestral allele. Despite having two outgroup species, we still observed an excess of fixed derived alleles in our data, likely due to ancestral allele miscalls. Nonetheless, the *B_2_* statistic modelled using minor allele frequency (MAF) spectra produced highly similar results, with the majority of outlier windows identified under MAF also detected in the DAF results. This reinforces the robustness of our candidate regions, but also highlights the potential of using MAF-based approaches for identifying balancing selection when reliable inference of ancestral alleles can be difficult (114), especially for non-models.

When comparing *B_2_* to *β*, we found limited overlap in top-scoring windows across populations; an unexpected result given that both methods aim to detect similar signatures of balancing selection. This discordance did not appear to differ substantially among populations. However, many of our most repeatable candidate regions showed elevated *β* scores, emphasising the value of integrating multiple populations and applying rank-based approaches (like PicMin; (75)) instead of relying on arbitrary thresholds. Finally, we compared our candidates to traditional measures of excess diversity (Tajima’s *D* and π). While top *B_2_* outliers typically ranked highly in these measures, this was not always true. This is likely due to the challenges in modelling the allele frequency spectra in small populations, where the assumptions underlying statistics like Tajima’s *D* do not always hold (115). Additionally, reduced linkage in certain populations, such as EU, may have lowered our overall power to detect balancing selection. However, by integrating information from all these complementary approaches, we narrowed our results to a high-confidence set of 23 candidate regions for functional investigation.

In summary, our study provides a comparative analysis of genome-wide scans for detecting balancing selection across multiple natural populations, revealing how NFDS may contribute to the maintenance of genetic diversity, even across demographically diverse populations. To our knowledge, this is among the first studies to explore evolutionary repeatability in the context of balancing, as opposed to positive selection, across natural populations. Our findings demonstrate the value of incorporating genome scans for balancing selection to complement phenotypic, ecological, and behavioural studies, offering a broader perspective on how NFDS operates. Future studies expanding genome-wide investigations of NFDS to additional taxa and demographic contexts will be critical to increasing our understanding of how balancing selection operates in maintaining diversity.

## Data Accessibility

All scripts pertaining to data analysis are available on GitHub (https://github.com/josieparis/guppy_balancing_selection).

## Author contributions

Josephine R. Paris - Formal analysis, Investigation, Methodology, Visualization, Writing - original draft, Writing - review & editing

James R Whiting - Formal analysis, Methodology, Writing - review & editing

Joan Ferrer Obiol - Formal analysis, Methodology, Visualization, Writing - review & editing

Kimberly A Hughes - Conceptualization, Funding acquisition, Project administration, Writing - review & editing

Bonnie A Fraser - Conceptualization, Formal analysis, Funding acquisition, Investigation, Methodology, Project administration, Writing - original draft, Writing - review & editing

## Acknowledgements

The authors wish to thank Helen Rodd for the collection of guppies from the Paria and Turure rivers and useful discussions at the conception of this project. Thanks to Mijke van der Zee for useful discussions and suggestions on analyses. Computational infrastructure support was provided by The University of Exeter’s High-Performance Computing (HPC) facility (ISCA). DNA sequencing was performed by the Earlham Institute, UK.

