## Supplementary Materials for "Repeated signatures of balancing selection in small and large populations of guppies (*Poecilia reticulata*)"

This supplementary document contains:

- Supplementary Figures S1 - S13
- Supplementary Methods 1
- Supplementary Methods 2
- Supplementary Methods 3

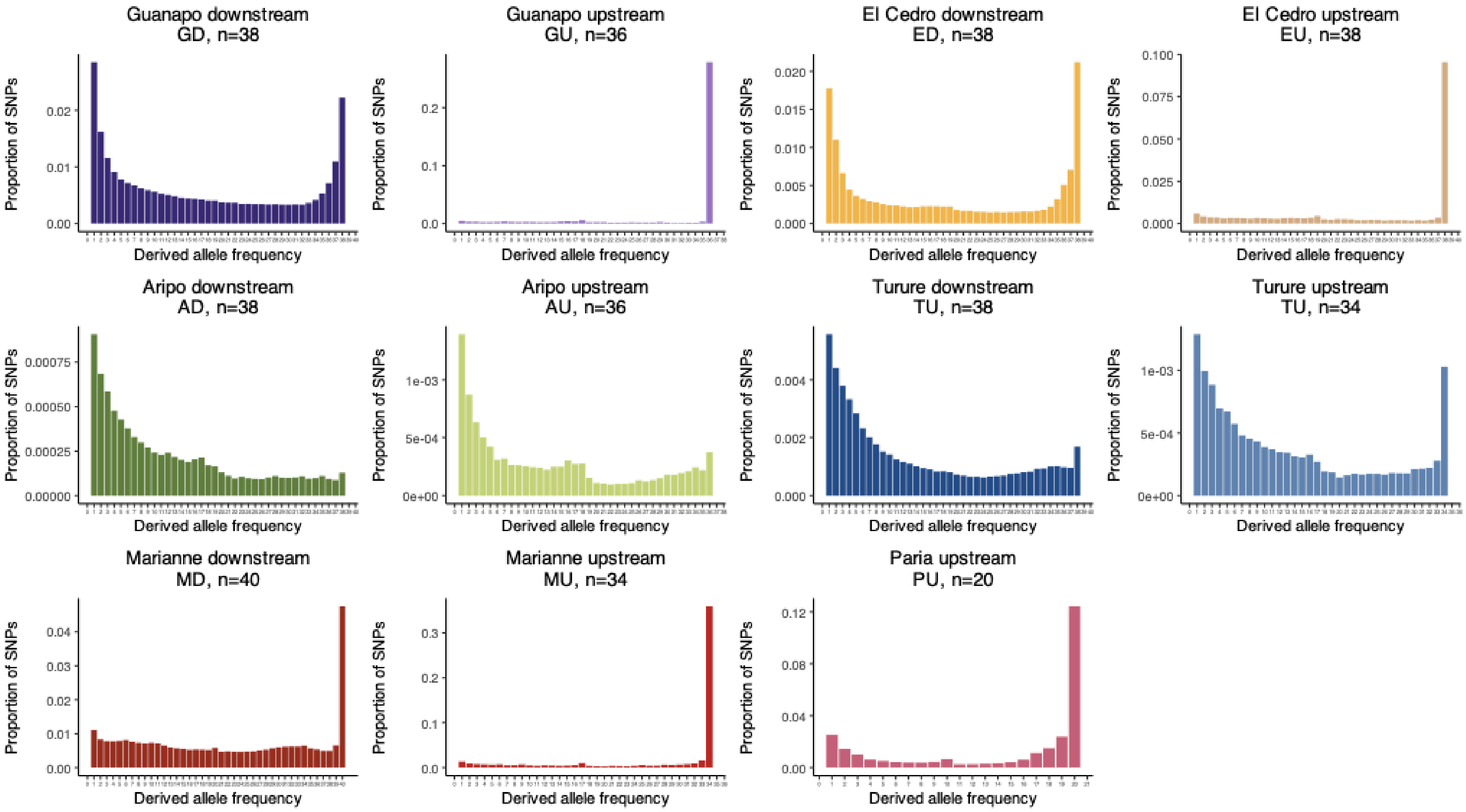


**Figure S1**. Plots of the derived allele frequency (DAF) spectra for each of the 11 populations. Populations are colour-coded as in the rest of the manuscript and supplementary figures.

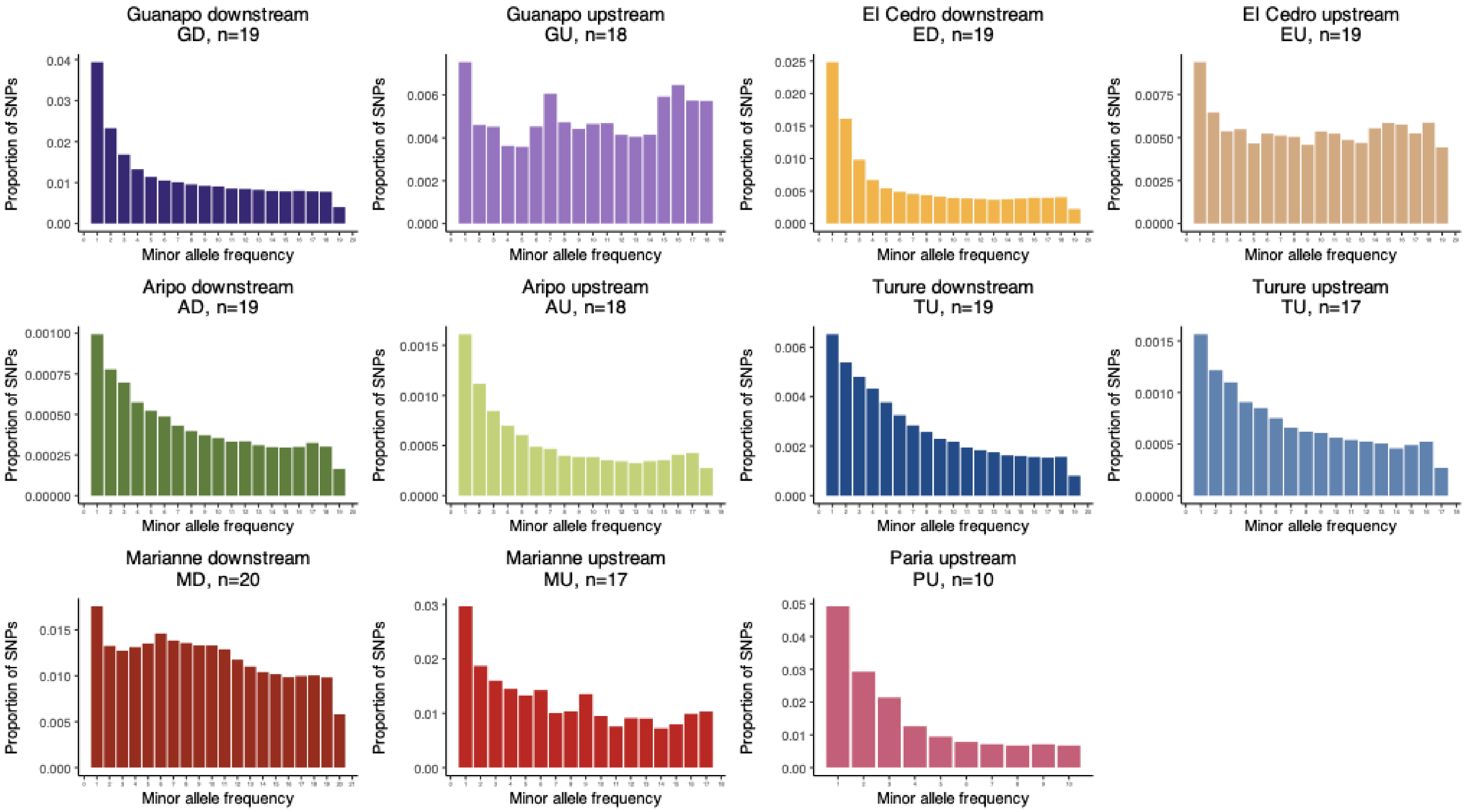


**Figure S2**. Plots of the minor allele frequency (MAF) spectra for each of the 11 populations. Populations are colour-coded as in the rest of the manuscript and supplementary figures.

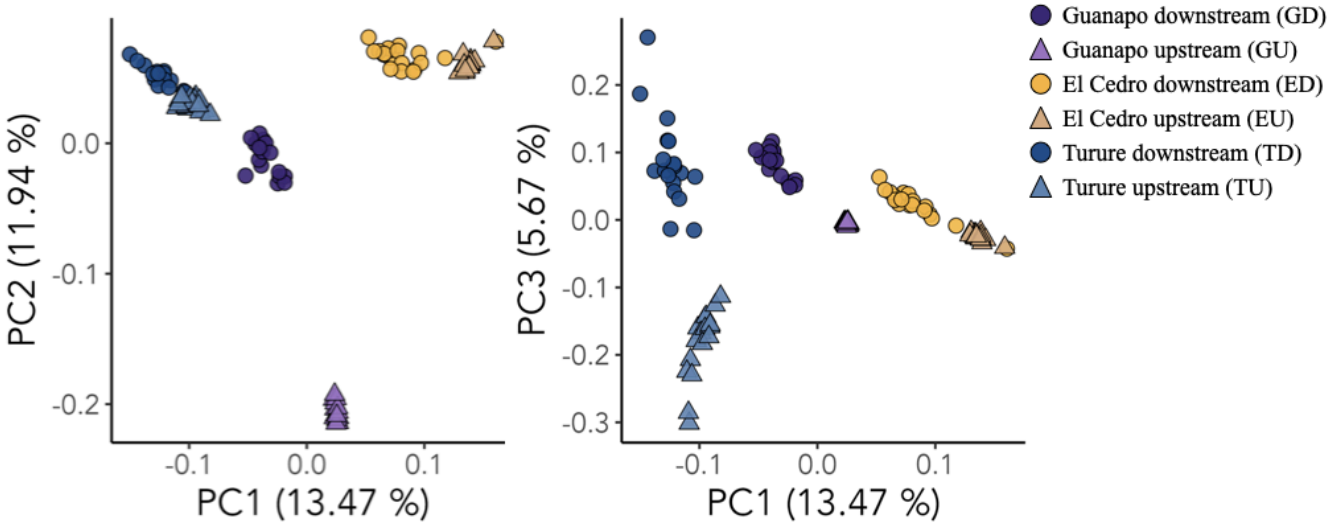


**Figure S3**. Principal Components Analysis (PCA) of the six populations showing Caroni drainage signatures. Populations include the natural upstream-downstream pair in Guanapo (GD-GU), the El Cedro populations (ED, EU), and the Turure populations (TD-TU). PC1 (13.47%) separates the El Cedro populations from the Guanapo populations (and the Guanapo-related introduction populations). PC2 (~12%) separates the Guanapo populations. PC3 (~6%) separates the TU introduction (introduced from GD) from its natural downstream counterpart (TD).

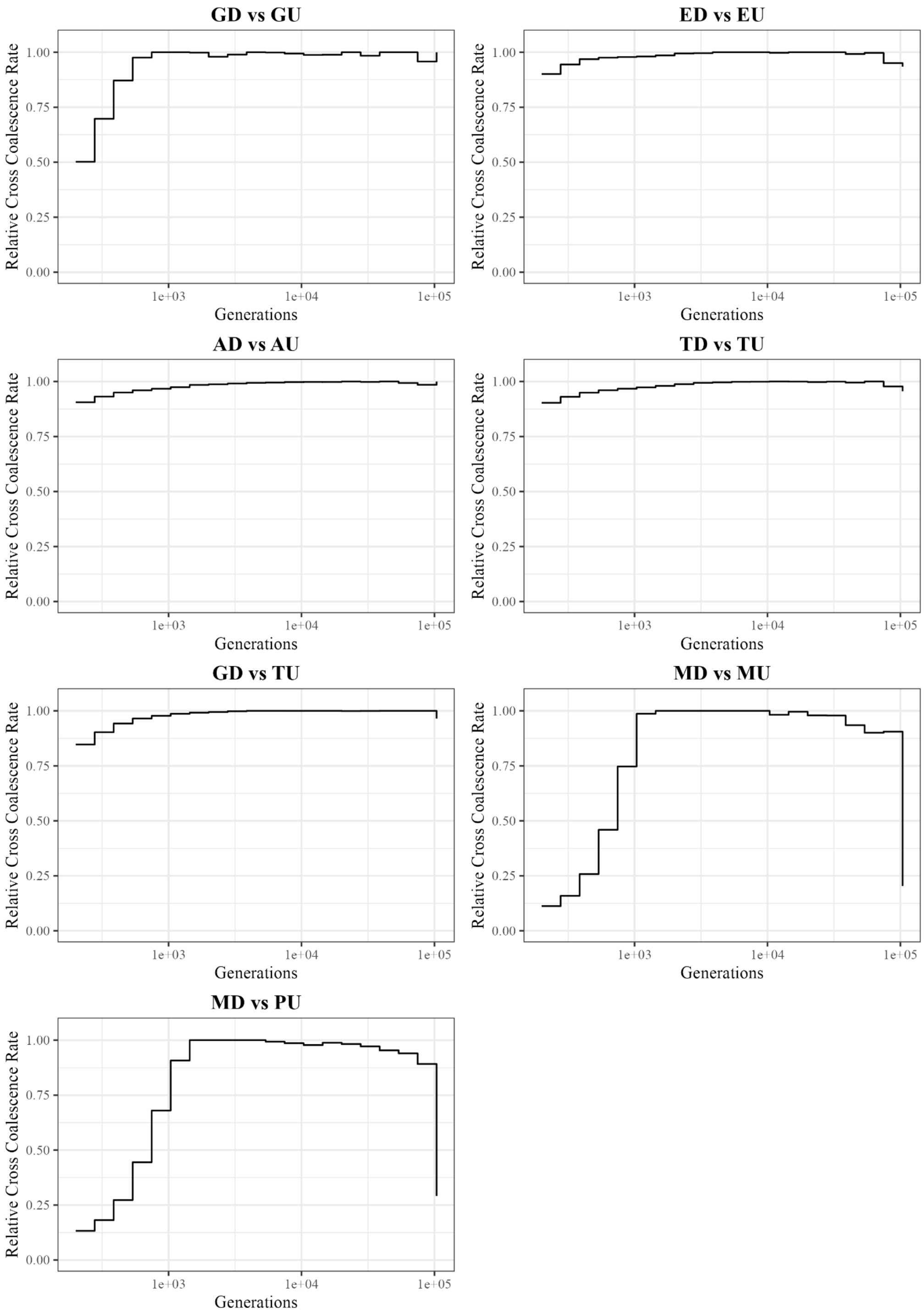


**Figure S4**. The relative cross coalescence rates (rCCR) calculated using Relate. Panels show the rCCRs calculated between: 1) natural downstream and upstream pairs: GD-GU; AD-AU; TD-TU; MD-MU; 2) experimentally introduced pairs: ED-EU; GD-TU, and 3) MD-PU (as PU and MU are shown have little population structure.

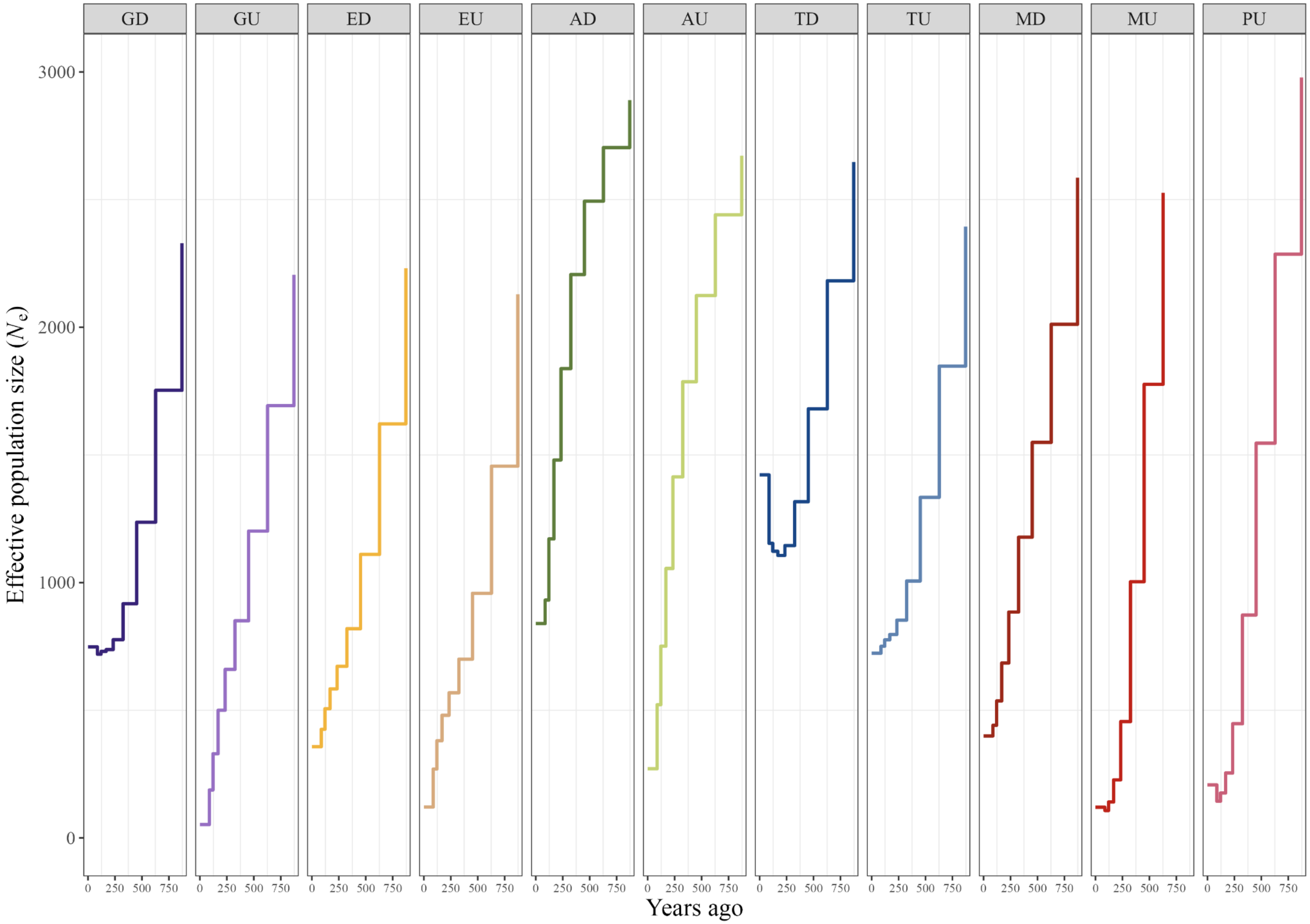


**Figure S5**. Recent changes in the effective population size ($N_e$) for the 11 populations calculated using Relate. Only the last 1000 years are shown to demonstrate the most recent demographic changes. Populations are colour-coded as in the rest of the manuscript and supplementary figures.

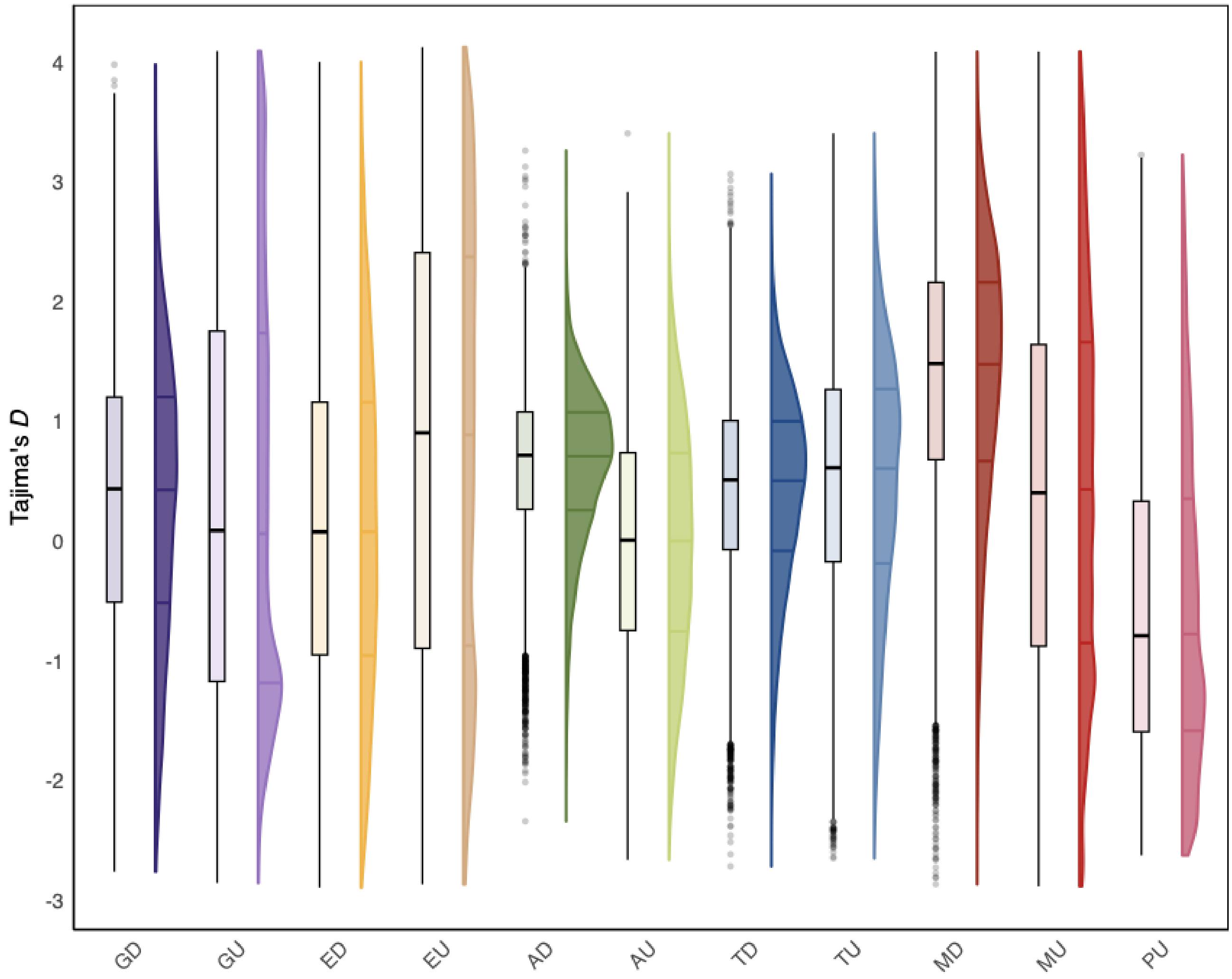


**Figure S6.** Raincloud plots showing the variation in Tajima's *D* (calculated in 50 Kb windows across the genome) for the 11 populations. Populations are colour-coded as in the rest of the manuscript and supplementary figures.

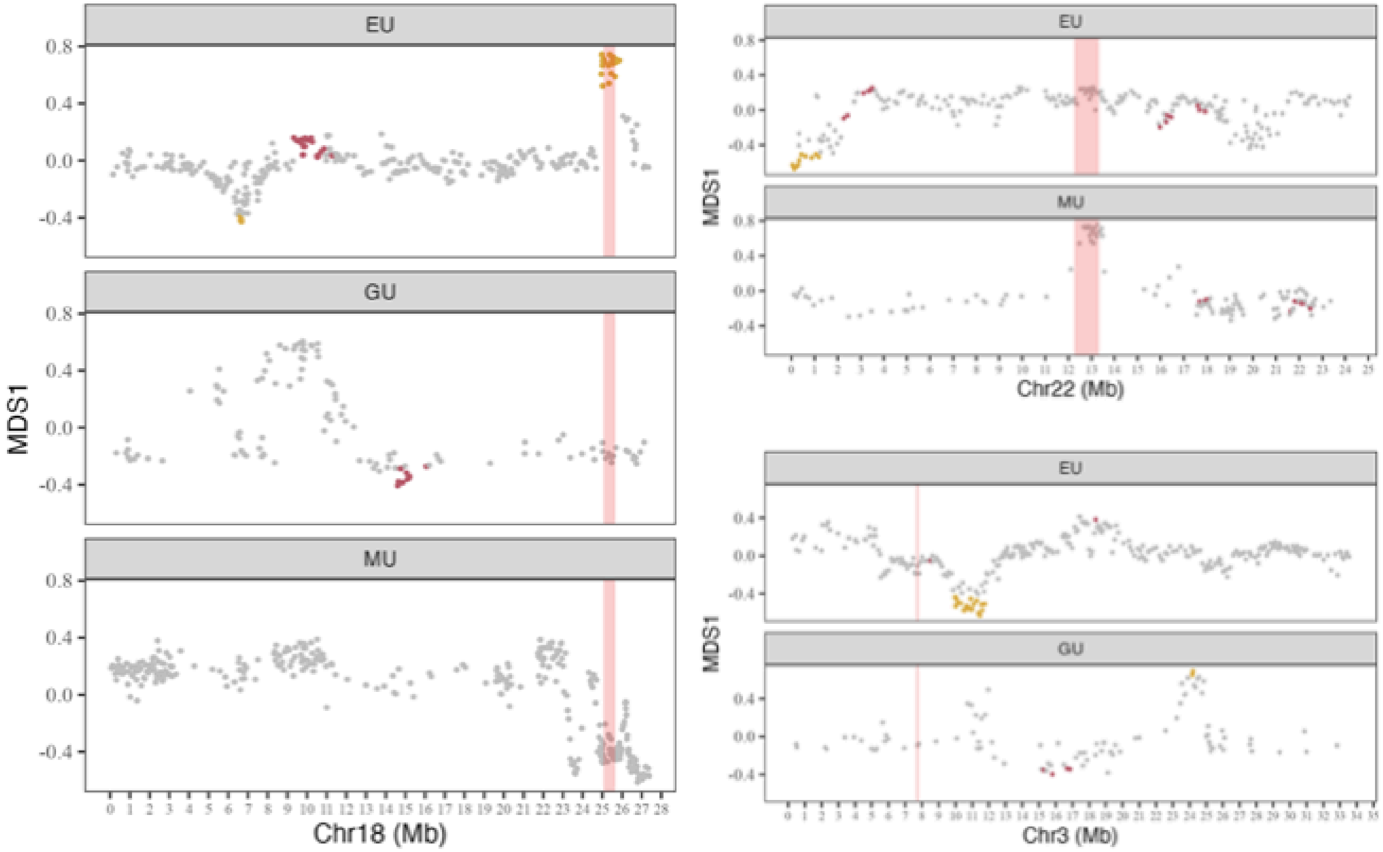


**Figure S7**. Local PCAs constructed using Lostruct for the three regions of overlapping elevated Tajima's *D* identified using change point detection (CPD) analyses: LG18, LG22 and LG3. Y-axis shows the MDS1 scores. Points are coloured by non-significant (grey), MDS1 significant (yellow) and MDS2 significant (red). Red highlighted regions in each plot represent the CPD identified region for each chromosome: LG18 (25.05 - 25.65 Mb) identified in EU, GU and MU; LG22 (12.25 - 13.35 Mb) identified in EU and MU; and LG3 (7.65 - 7.80 Mb) identified in EU and GU.

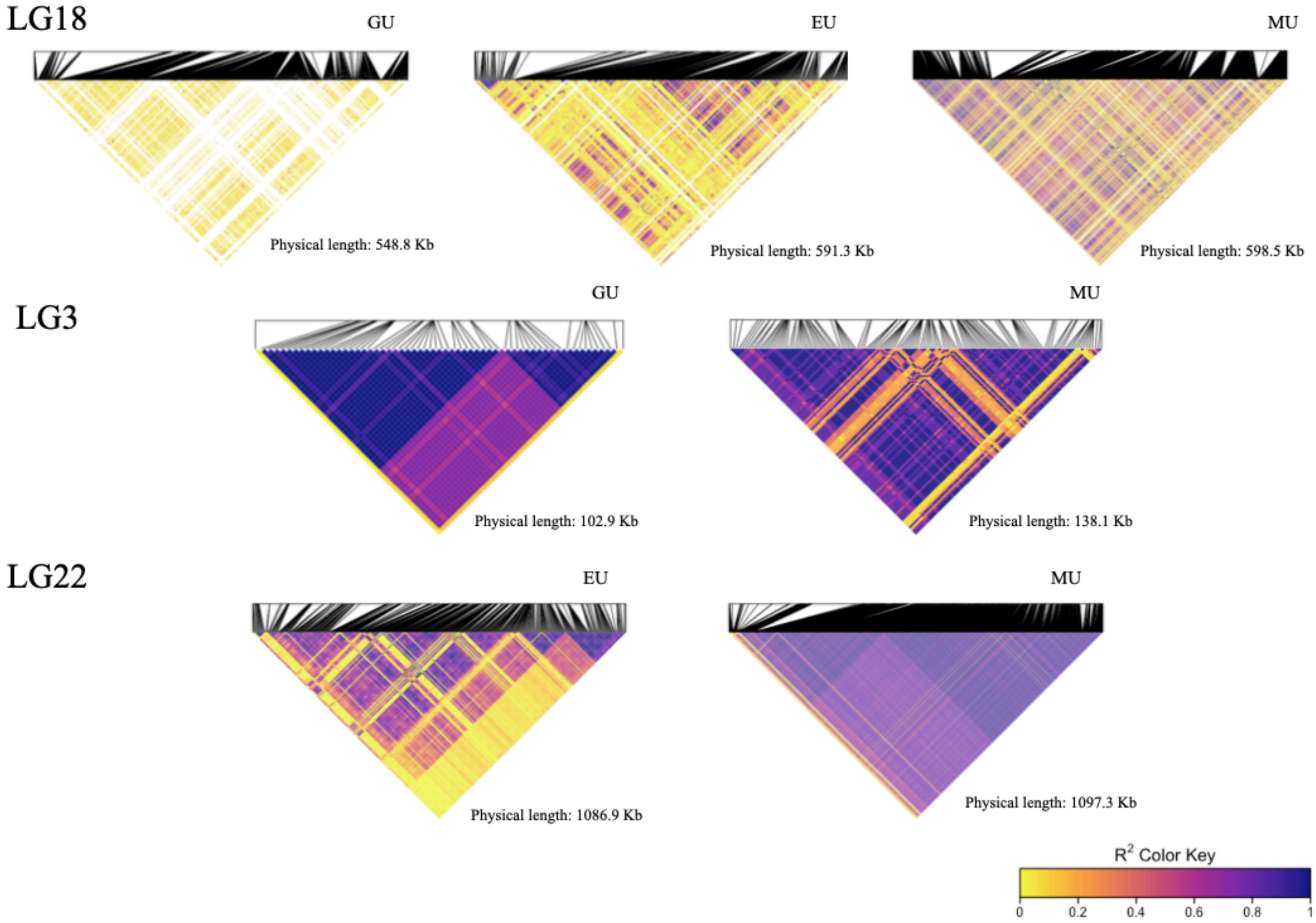


**Figure S8**. Patterns of Linkage Disequilibrium (LD) calculated in the small populations identified as showing overlapping change point detection (CPD) regions of elevated Tajima's *D*: LG18 (25.05 - 25.65 Mb) identified in EU, GU and MU; LG3 (7.65 - 7.80 Mb) identified in EU and GU, and LG22 (12.25 - 13.35 Mb) identified in EU and MU.

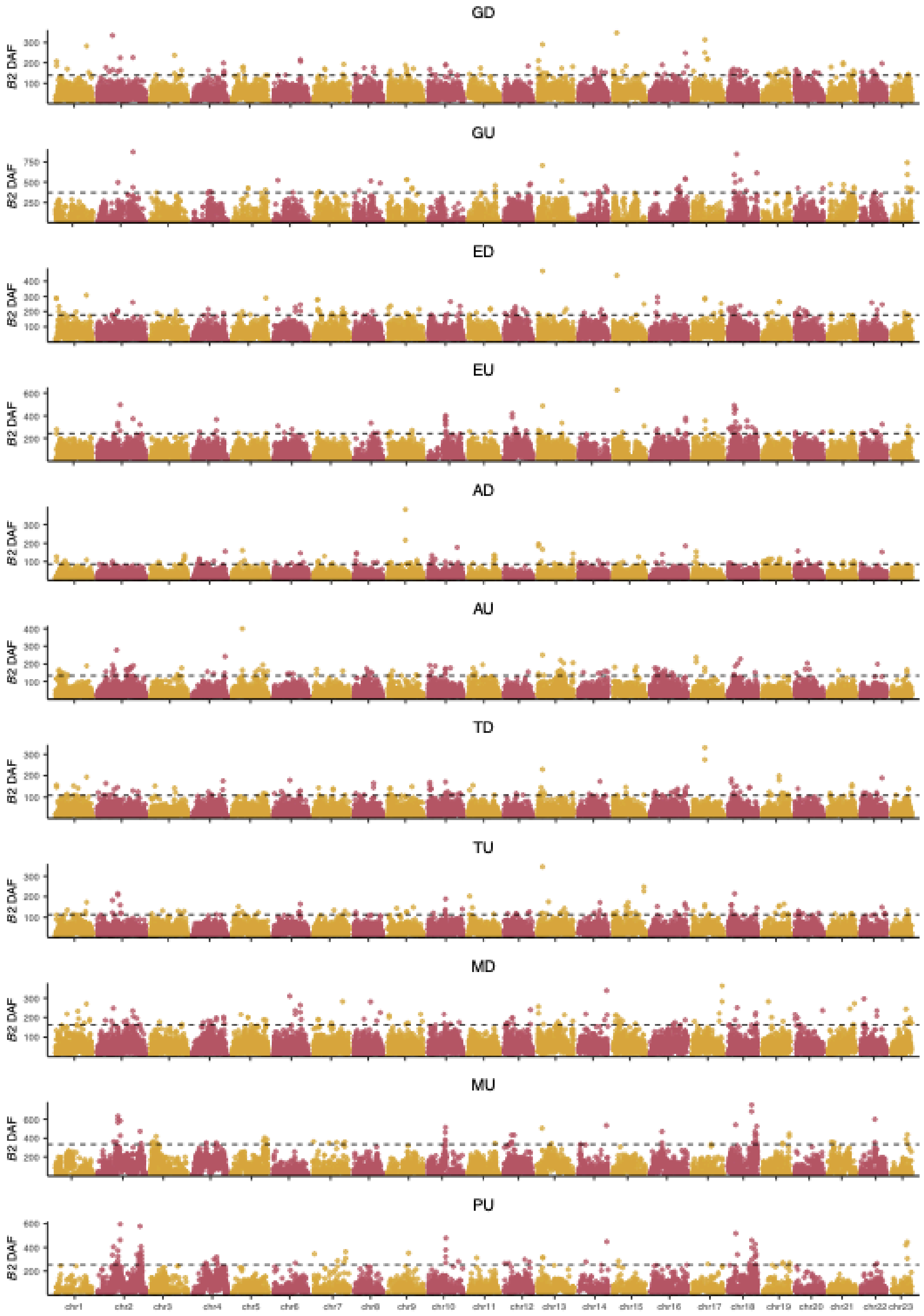


**Figure S9**. Manhattan plots showing the $B_2$ CLR scores calculated using the derived allele frequency (DAF) spectra across the 23 chromosomes of the guppy genome for the 11 populations. Black dashed lines show the upper 99% cut-off (*i.e.*, top 1%) $B_2$ distribution for each population.

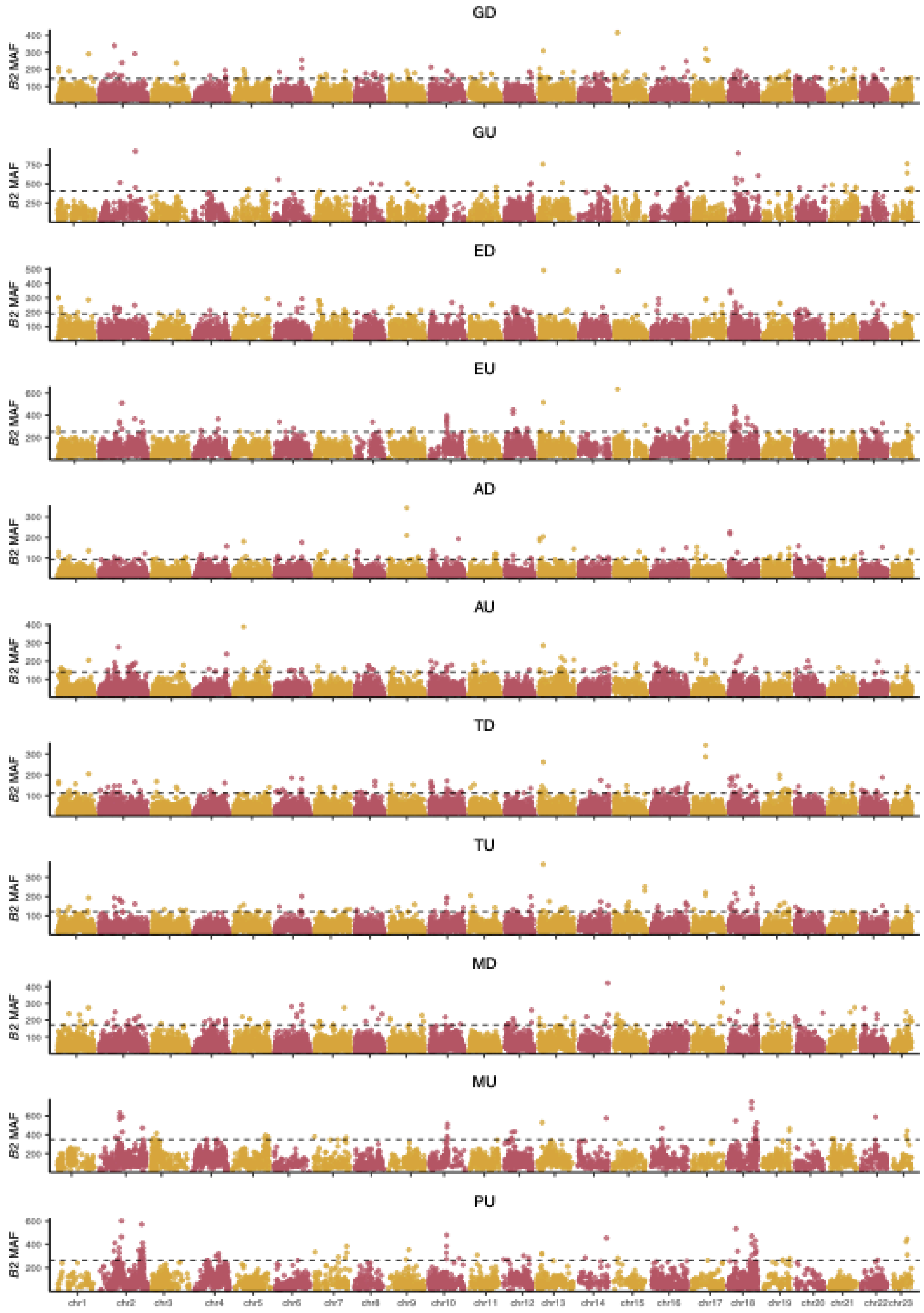


**Figure S10**. Manhattan plots showing the $B_2$ CLR scores calculated using the minor allele frequency (MAF) spectra across the 23 chromosomes of the guppy genome for the 11 populations. Black dashed lines show the upper 99% cut-off (*i.e.*, top 1%) $B_2$ distribution for each population.

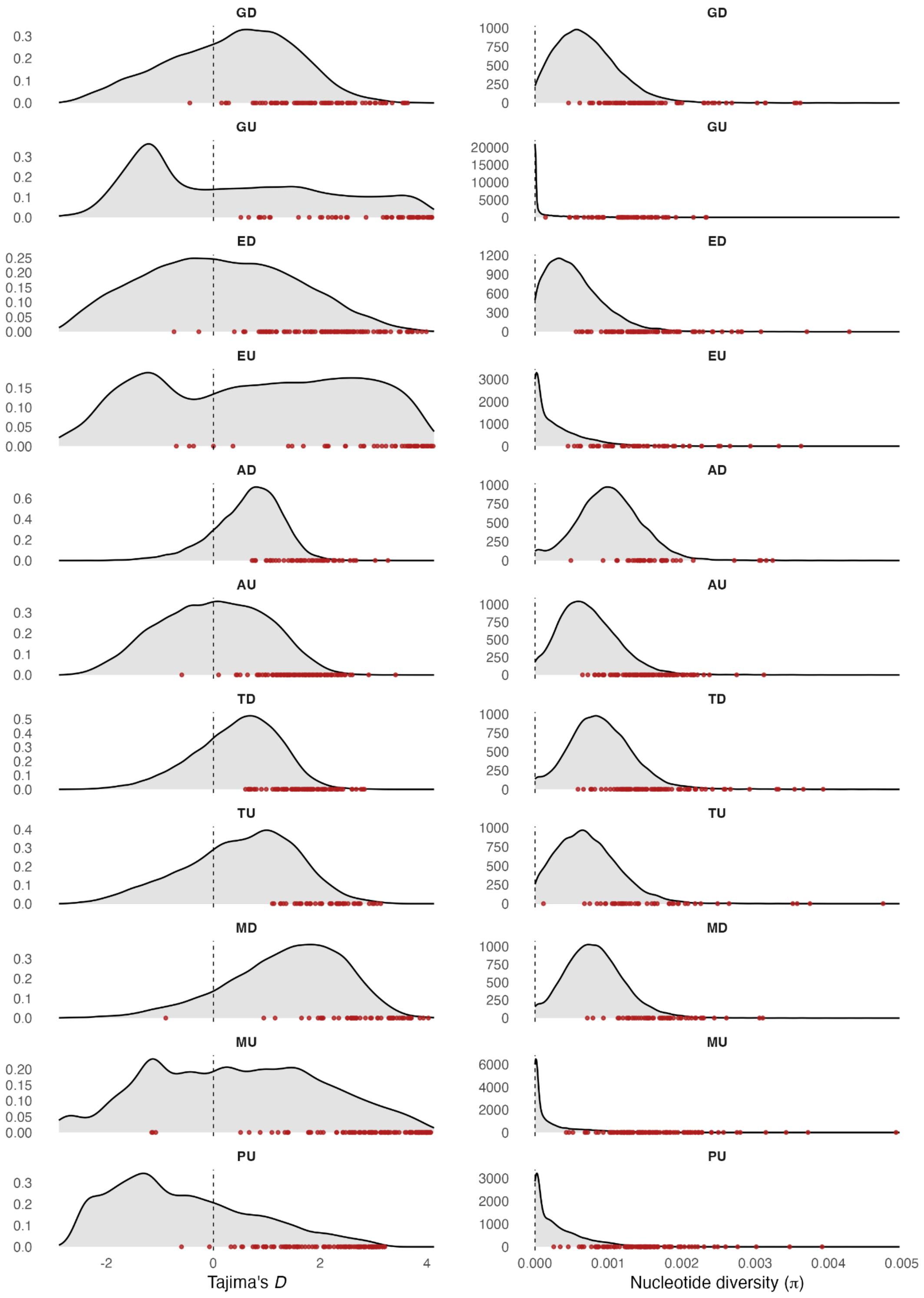


**Figure S11**. Plots showing the distribution of Tajima's *D* (left) and nucleotide diversity, π (right) for each of the 11 populations. Both statistics were calculated in 50 Kb windows. Red points mark the significant $B_2$ windows (>99%, ŝ ≥ 9).

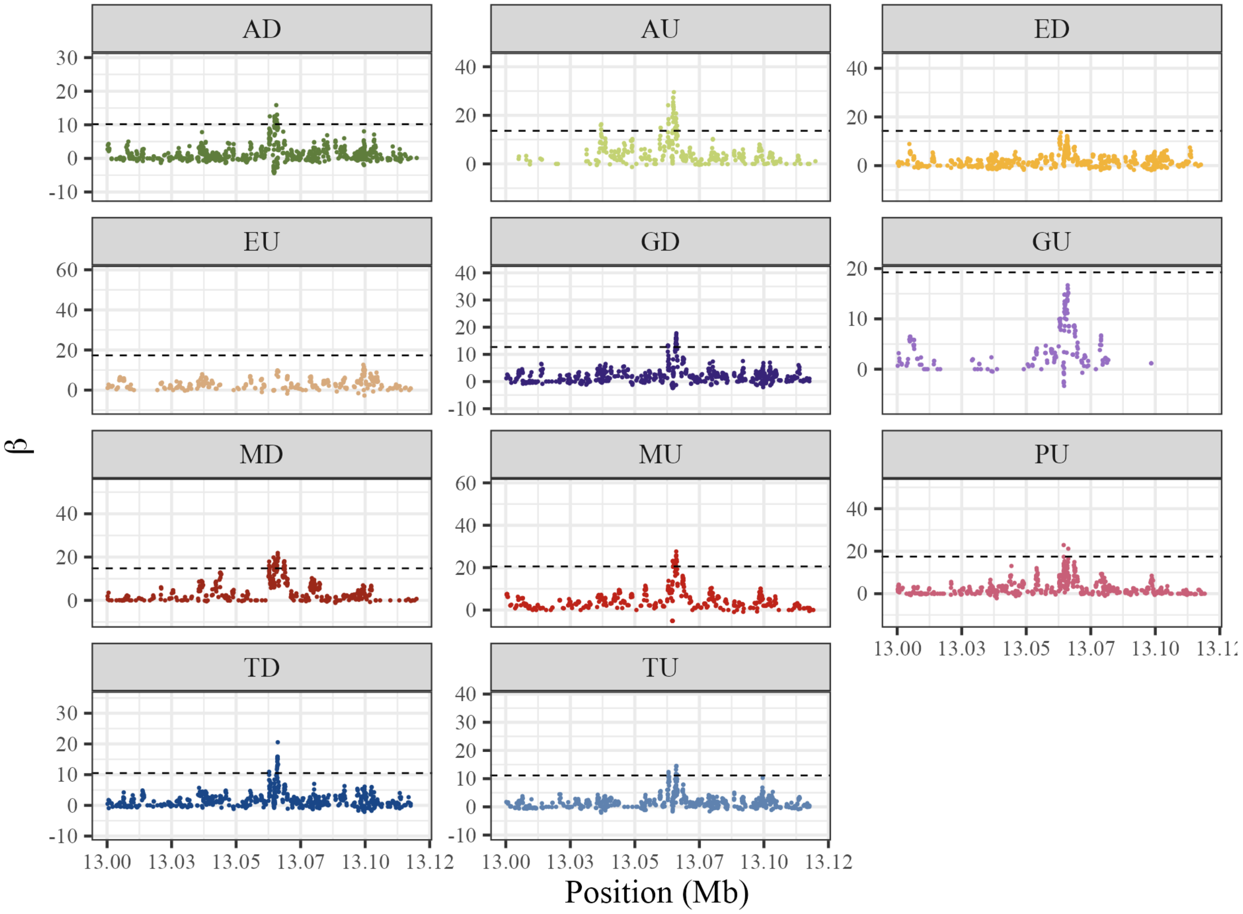


**Figure S12**. Per-SNP β scores for the candidate region on LG22 across the 11 populations. Populations are colour-coded as in the rest of the manuscript and supplementary figures. The dashed horizontal line represents the 99% threshold (*i.e.*, top 1%).

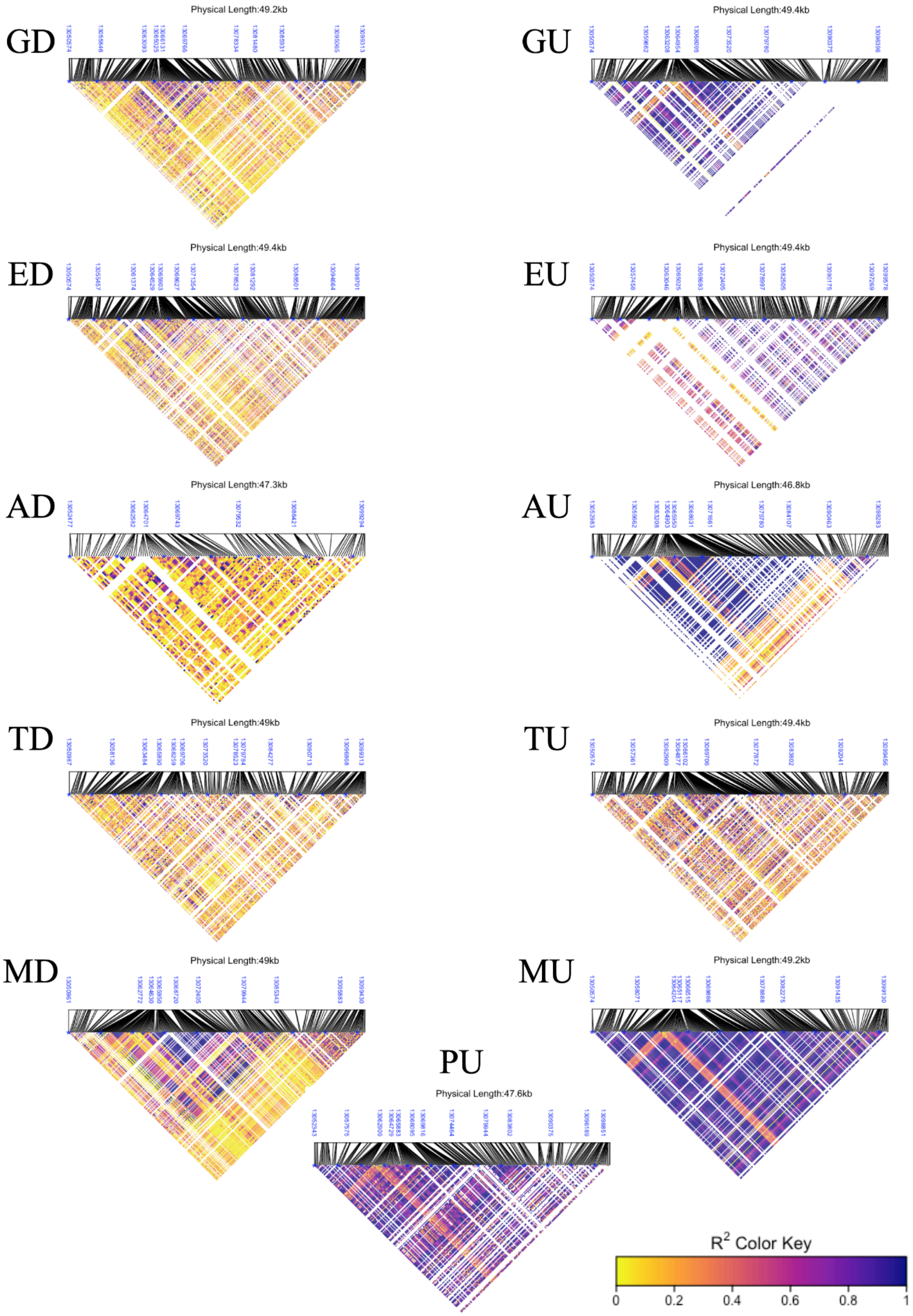


**Figure S13**. Linkage Disequilibrium (LD) calculated in the LG22 candidate region (13.05 - 13.10 Mb) across the 11 populations.

### Supplementary Methods

#### Methods S1: Divergence dating

For estimating divergence dates between the 11 populations we included data from 36 individuals: three individuals (with the lowest amount of missing data) from each of the 11 populations and three individuals from the sister species, *P. wingei*. To enable the correct calling of monomorphic sites in the sister species, individuals were joint-genotyped together in GATK and filtered (as in the main methods section) (Table S3). To calibrate the tree, we followed the recommendations of Stange et al. (2018) by constraining the root of the tree (divergence between *P. reticulata* and *P. wingei*) to follow a normal distribution with a mean of 3.41 mya and a standard deviation of 0.329 (Whiting et al., 2021). We performed three runs of 1,000,000 MCMC iterations sampling every 1000 iterations with each run using a different set of 3000 unlinked SNPs (minimum distance between SNPs = 10 Kb). Convergence and stationarity of the estimated parameters were confirmed for each run (effective sample sizes > 300) using Tracer v1.7.1 (Rambaut et al., 2018), discarding the first 10% of MCMC iterations as burn-in. We used Tracer to check that the three runs converged towards the same values for the following parameters: speciation rate, age of the root of the tree, and substitution rate. Trees sampled from the posterior were visualised using DensiTree v.2.2.7 (Bouckaert, 2010).

#### Methods S2: Designation of the derived allele

To make use of the derived allele, *P. reticulata* variants were jointly-genotyped alongside the *P. picta* and *P. wingei* data in GATK4 v4.1.8.1 (McKenna et al., 2010), following the filtering approaches outlined in the Main Methods. The ancestral allele was called as a consensus between *P. reticulata*, *P. picta* and *P. wingei* and was written as the ALT allele in a draft VCF file using a custom python script, which resulted in the flipping of 8.4% of the REF alleles in *P. reticulata* and an overall SNP recall rate of 94%. The ALT allele was then written to an ancestral fasta file using bcftools v1.9 consensus (Danecek et al., 2021), which was used to populate the ancestral-allele (i.e., AA) column using fill-aa in VCFtools (Danecek et al., 2011). Further details and scripts can be found on the GitHub repository: https://github.com/josieparis/guppy_balancing_selection

#### Methods S3: VCF filtering prior to scans of balancing selection

To remove artefacts which can confound scans for balancing selection, variants were filtered prior to conducting scans for balancing selection. The original VCFs were filtered for regions with 1) low-mappability; 2) high repeat content and 3) excess low or high coverage variants.

Mappability was computed over a range of *k*-mer sizes (100, 125, 150, 200) using GenMap (Pockrandt et al., 2020). Intersect between mappability called across the different *k*-mer sizes was assessed in R, selecting regions of reduced mappability where three or more kmer sizes overlapped using bedtools v2.27.1 (Quinlan & Hall, 2010). Bedtools *maskfasta* was used to create a positive mappability mask bed file, which was intersected with the VCF file using bedtools *intersect*.

Repetitive regions were calculated across 10 Kb windows of the genome using bedtools and the resulting distribution was plotted in R. We considered a window to be repetitive when repeats comprised > 15% of the window. bedtools *intersect* was used to remove highly-repetitive windows.

Coverage was calculated using vcftools v0.1.6 (Danecek et al., 2011) --site-mean-depth and the distribution was plotted in R. The VCF file was then filtered using a site-min depth of 7 and a max-mean-depth of 20 in vcftools.

Further details can be found on the GitHub repository: https://github.com/josieparis/guppy_balancing_selection

#### Methods S4: Re-annotating the guppy genome

The currently available annotation for the guppy reference genome GCA_904066995 (Fraser et al., 2020) was created using BRAKER2 (Brůna et al., 2021) using only single-end RNA-seq data. We took the opportunity to re-annotate the genome using BRAKER3 (Gabriel et al., 2024; Stanke et al., 2006, 2008), which makes use of both raw RNA-seq data and a protein database.

To generate evidence for BRAKER3, we used HISAT2 v2.2.1 (Kim et al., 2019) to map a multi-tissue mRNA-seq dataset consisting of fin (SRR10521935), ovary (SRR1137868), testes (SRR1140963), and a mixed organ pool from female (SRR1045559) and male (SRR1045560) guppies to the soft-masked genome. Protein data included the Vertebrata partition of OrthoDB v11 (Kuznetsov et al., 2023) and the proteomes of six Poecilidae species (*Poecilia reticulata*, *Poecilia formosa, Poecilia latipinna, Poecilia mexicana, Xiphophorus couchianus, Xiphophorus maculatus*), downloaded from the Ensembl Release 113 (Harrison et al., 2024). AGAT v0.7.0 (Dainat, 2024) was used to provide basic filtering for structural anomalies and to quantify statistics regarding structural aspects of the protein-coding regions.

For functional annotation, we performed a search of each predicted protein sequence against the InterPro protein database using InterProScan v5.72-103 (Blum et al., 2021; Jones et al., 2014) and with eggNOG-mapper v2 (Cantalapiedra et al., 2021) against the eggNOG v5 orthology database (Huerta-Cepas et al., 2019). Quantitative and qualitative assessment of the gene predictions and functional annotation was performed at every step of the annotation process using AGAT v0.7.0, BUSCO v5.7.1 (Manni et al., 2021; Simão et al., 2015) against the actinopterygii_odb10 dataset (n=3640) from OrthoDb v10 (Kriventseva et al., 2019), and OMArk v0.3.0, using OMAmer v2.0.3 (Nevers et al., 2024) with the Cyprinodontoidei clade (n = 18451).

The final annotation had a BUSCO score of 94.5% complete (92.7% as single-copy, 1.8% as duplicated), with 2.6% fragmented, and 2.9% missing and an OMArk score of 97.19%

complete (83.59% as single copy, 13.6% as duplicated), with 2.81% missing. OMArk also showed a consistent lineage placement of 64.91% and no contamination. Functional annotation provided gene names to 22,873 genes, and gene descriptions to 31,669 genes. All annotation files are available at the GitHub repository: https://github.com/josieparis/guppy_balancing_selection
